# Geometric characteristics of cubically symmetric and triply periodic scaffolds for optimal cell migration

**DOI:** 10.64898/2026.04.13.718106

**Authors:** C. Lonati, L. Preziosi

**Affiliations:** Dipartimento di Scienze Matematiche “Giuseppe Luigi Lagrange”, Politecnico di Torino, Italy

## Abstract

In tissue engineering, it is important to conceive and construct artificial bio-mimetic scaffolds able to foster cell migration as this is a fundamental process in wound healing and tissue regeneration. In order to do that, cubically symmetric and triply periodic porous structures have been identified as promising candidates for instance for the reconstruction of artificial cartilages and bones, also due to their tunable mechanical characteristics and highly inter-connected porous architectures that mimic the trabecular bone hyperboloidal topography. We propose here a mathematical approach that might be helpful to identify what are the best geometrical characteristics of such scaffolds, in order to promote cell migration into the porous structures and speed-up their re-population. The method is based on the observation that cell nucleus deformations should be avoided, yet assuring a good possibility for the cell to reach the wall of the porous structure. Mathematically speaking, this leads to the problem of identifying the size of the largest sphere that can pass, without being stuck, through the pores of the bio-mimetic scaffold.

## 1 Introduction

In the human body, cell migration in the three-dimensional network of fibers that make up the extracellular matrix (ECM) of many tissues plays a crucial role in many physiological and pathological situations. Notably for the purpose of this article, in wound healing processes, cell migration is fundamental to repair both basement membrane-underlaid epithelia and connective tissues [26, 29]. So, it is clear that the faster the process is, the better it is. Having this purpose in mind, in order to reconstruct tissues in vitro as fast as possible, a crucial direction of tissue engineering is the identification of proper bio-mimetic structures with optimal characteristics favoring cell migration and proliferation.

The key factors governing cell migration in both two- and three-dimensional environments are the dynamic adhesion of cells via the expression of trans-membrane proteins (mainly integrins) attaching to the ECM and the generation of the traction forces necessary for advancement by contraction of cytoskeletal elements (see, for instance, [14]). However, in order to migrate within three-dimensional porous environments, cells also need to find their way through steric obstacles, constituted, for instance, by the network of ECM fibers themselves, that can hamper their migration [6, 21, 28]. In such situations, the cell nucleus, which is the stiffest organelle of the cell, must deform considerably and strong cytoskeletal forces must be exerted to pass through restricted openings in the ECM. Actually, in extreme cases, environmental conditions are so harsh to represent a real physical limit of migration and the only way a cell can migrate is by activating proteolysis to degrade ECM fibers and open the space for cell movement [1, 7, 8, 13, 22]. In any case, such situations should be avoided in an artificial scaffold because they slow down cell migration.

From the modeling point of view, several articles have studied how migration is affected by the mechanical characteristics of the cells and of the ECM and by the geometrical characteristics of the environment with respect to the size of the cell and more importantly of its nucleus. For example, in [9] and [10] a continuum mechanics method is used to evaluate the energy required to permit the entrance of the nucleus into a microchannel. This method allowed to identify a criterium of penetration in microchannels that depends on one hand on a dimensionless number measuring the traction ability of the cell versus the stiffness of the nucleus and on the other hand on the ratio of the diameters of the nucleus with respect to size of the microchannel. Scianna and coworkers [23, 24, 25] used a cellular Potts model to describe cell migration in microchannels, ECM, or arrays of micropillars. The last geometrical set-up was also used in [4, 17]. In particular, Vermolen et al. [4] propose a two-dimensional discretized model of both the cellular and nuclear membrane. Finally, Giverso et al. [11] developed a single cell-based model that includes the mechanical properties of both the cellular and nuclear membrane, the asymmetrical front-back renewal of the actin cytoskeleton, and the contact interactions between the cell and the external environment. The migration through a corrugated channel was then studied.

In addition to the identification of criteria describing the physical limits of migration, the models in [11, 23, 24, 25] showed that the maximum speed of migration is achieved when the channel size is close to the diameter of the cell nucleus. In fact, as can be easily understood, if the characteristic pore size is slightly larger than the nucleus diameter, the cell encounters little impediment to migrate into the porous structure. For smaller sizes, some work needs to be done by the cell to squeeze its nucleus into the pore, which slows down migration. At the other extreme, if the pore size gets too large, then the cell will encounter difficulties in adhering to the pore wall and pulling in an effective way. Overall, these phenomena give rise to a bimodal behavior that is actually observed in experiments. For example, bimodal relationships between cell migration ability and the size and deformability of 3D matrix scaffolds have been observed in smooth muscle cells migrating in polyacrylamide substrates with tunable mechanical properties [19], and in mouse fibroblasts cultured in stepwise cross-linked collagen-glycosaminoglycan matrices of constant pore size [12]. A bimodal dependence on matrix rigidity was also reported in isotropic homogeneous networks, as in the case of prostate cancer cells embedded in Matrigel with a fixed fibronectin level and variable stiffness [32].

The knowledge of the mechanisms discussed above can be exploited in tissue engineering to design synthetic implant materials, i.e., artificial scaffolds, trying to optimize their pore size and stiffness to accelerate cell migration and in-growth [2]. This would be particularly relevant for regenerative treatments and tissue reconstruction, notably dermal layers, tendons, cartilages, and bones. Specifically, we focus on some structures that are found to be particularly suitable for cell attachment, migration, and proliferation and for the necessary transport of nutrients inside the porous structure, namely the offset cubic lattice and some triply-periodic structures (e.g., Schwarz surfaces and gyroids) [3, 20, 30, 31]. In fact, these scaffolds are characterized by a high surface-to-volume ratio and porosity. For this reason, for instance, gyroids are also used for the construction of heat exchangers and for other systems in automotive and aerospace due to the great stiffness-to-mass ratio and high energy absorption. In addition, it has been found that the mechanical properties of such structures can be easily tuned so that in cartilage and bone repair the scaffold can optimally integrate in the surrounding tissue avoiding stress shielding [15]. The same flexibility in the construction of scaffolds with different geometrical structures and mechanical properties may, in principle, be tuned in a heterogeneous way to drive stem cell fate and differentiation in a space-dependent way [27]. Finally, the interconnected porous structure looks promising for the necessary vascularization of the tissue, that in three dimensional applications represents a problem to be properly addressed.

In this framework, the technological thrust motivating this article is the need to identify the best geometrical characteristics of some triply-periodic structures that can favor the process of cell migration in artificial scaffolds in order to foster its rapid re-population. The driving idea is then that during migration in the porous scaffold nucleus deformation needs be avoided. From the mathematical viewpoint this means to find the relationship between the size of the periodic cell of the suggested structure, that is usually given in implicit form in terms of trigonometric functions, and the diameter of the biggest sphere that can pass through its pores.

Thanks to the symmetry and periodicity of the structures studied, we are able to find analytical solutions determined using the following strategy: we look for contact points on the triply periodic surface that have directly opposite normals and then, among these, we look for the pair that has the minimum distance. Non-interpenetration of matter is also controlled, which translates in comparing the curvatures of the sphere and of the surface at contact points, guaranteeing local injectivity of the configurations [16].

We find, for example, that if *l* is the spatial periodicity of the structure and *d* is the diameter of the cell, in the case of a Schwarz P-surface 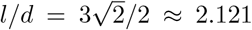, in the case of a Schwarz D-surface *l/d* ≈ 2.846, while for the gyroid 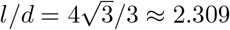.

The plan of the paper is as follows. In Section 2 we present the structures that will be analyzed and summarize the main results. In Section 3 we study cubic lattices and we explain the general method that can be used to treat triply periodic minimal surfaces. In the following sections (and appendices) we deal with the analysis of the Schwarz P-surface (Section 4), of the Schwarz D-surface (Section 5) and of the gyroid (Section 6). In all cases we give details that lead to results that are more intuitive for the Schwarz P-surface (which can then be considered as a test case) and less intuitive for the gyroid.

## 2 Geometrical Structures and Main Result

In the present work we will analyze the following structures shown in Table 1:

**Table 1:**
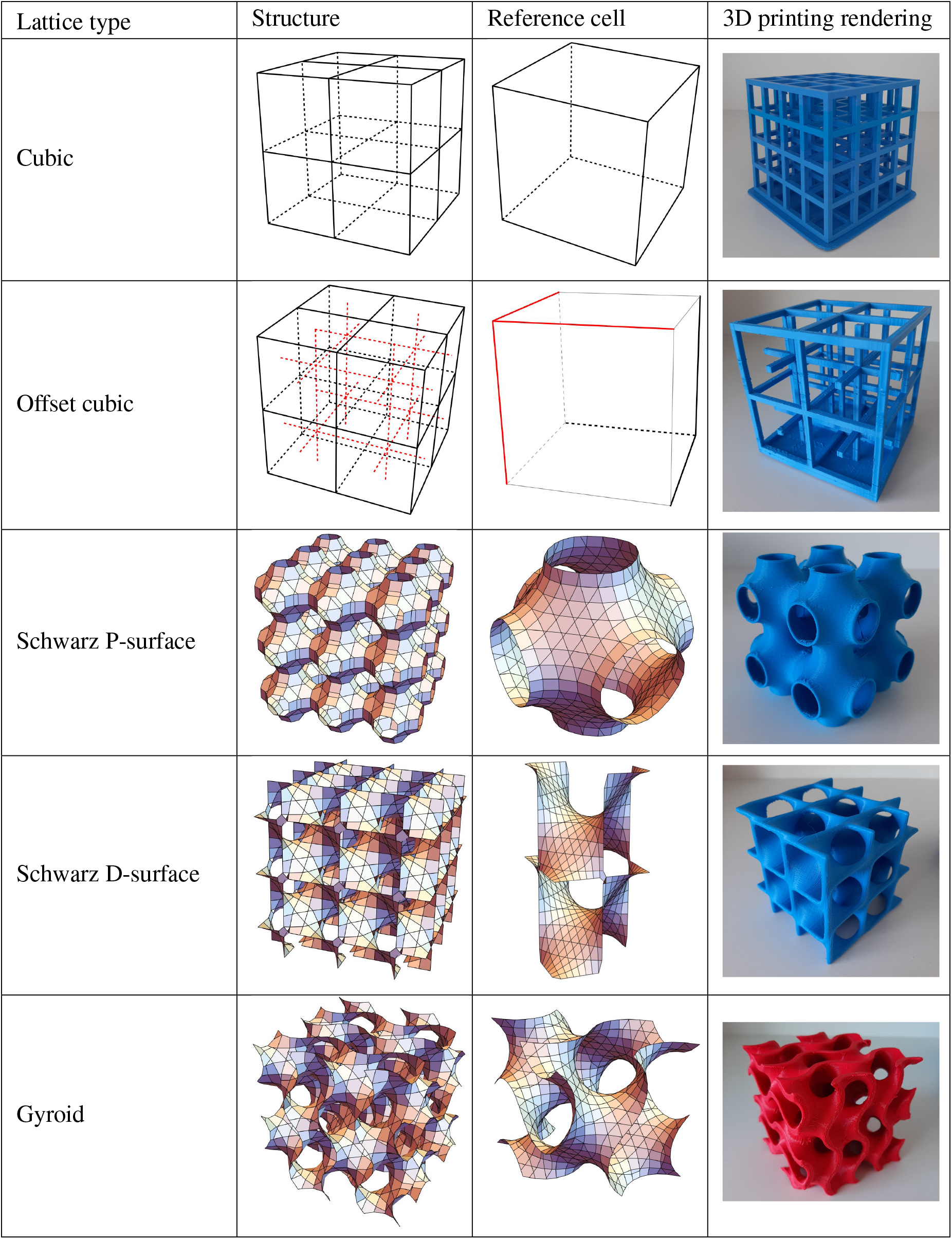
The five porous structures and their rendering through 3D printing.

- **Cubic lattice:** made by a regular array of cubes with the length of the edges equal to *l* (first row in Table 1);
- **Offset cubic lattice:** made by two interlaced cubic lattices that are ideally disconnected. It can be obtained as follows: taken a standard cubic lattice with the length of the edges equal to 2*l*, position an identical dual cubic lattice with vertices in the midpoints of the cells of the original lattice, i.e. shifted by (*l, l, l*) (second row in Table 1). The reference cell is a cube having a side equal to *l* (that is not periodic in space, but repeats with different rotations), with just 6 edges, arranged in two triples of perpendicular edges converging in two opposite vertices of the cube;
- **Schwarz P-surface:** triply-periodic minimal surface having the shape of an inflated cubic lattice separating the space in two intertwined congruent labyrinths. It is described by the dimensionless implicit relation cos *x* +cos *y* +cos *z* = 0. The periodic cell is a cube of edge 2*π*, and we will study it in the set (−*π, π*]^3^ (third row in Table 1);
- **Schwarz D-Surface:** triply-periodic minimal surface with two intertwined congruent labyrinths, described by the dimensionless implicit relation cos *x* cos *y* cos *z* − sin *x* sin *y* sin *z* = 0. The periodic cell is a cube of edge *π*, and for convenience we will study it in the set (−*π/*2, *π/*2]^2^ *×* (−*π, π*] (fourth row in Table 1);
- **Gyroid:** triply-periodic minimal surface described by the dimensionless implicit relation sin *x* cos *y* + sin *y* cos *z* + sin *z* cos *x* = 0. The reference cell is a cube of edge 2*π*, that for convenience we will take to be the set (− *π, π*]^3^ (fifth row in Table 1).

In reality, of course, from the manufacturing point of view it is clear that the segments of the cubic lattices and the surfaces have a finite thickness. This means, for example, that the region of space that is filled by the output of a 3D-printer is for instance given by |*F* (**x**)| *< ε* where *F* (**x**) is one of the implicit functions above and *ε* is of the order of the dimensionless thickness of the pore wall, i.e. scaled with the size of the reference cell. For example, for the P-surface the dimensional thickness of the wall is 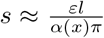 where 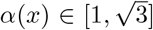 and *l* is the size of the periodic cell. However, in the following, for the sake of simplicity, we will work in the limit *ε* → 0.

While the symmetries characterizing the cubic lattices are quite clear, those characterizing triply-periodic surfaces might not be visually immediate. They can be better understood by looking at the symmetries of the implicit formulas defining them. It is then found, for instance, that

- the Schwarz P-surface is symmetrical with respect to the three coordinate planes (e.g., changing *x* in − *x*, that is a flip of the *x*-axis), to the three axes (e.g., changing *y* in − *y* and − *z* in *z*, that also represents a rotation of 180° around the *x*-axis), to the origin, and it is invariant for any permutation of the three coordinates;
- the Schwarz D-surface is symmetrical with respect to the three axes and invariant for any permutation of the three coordinates;
- the gyroid is symmetrical with respect to the origin, and invariant for any cyclic permutation of the three coordinates (e.g., changing {*x, y, z*} into {*y, z, x*}), that corresponds to a rotation.

The aim of the following sections will be to give an evaluation of the diameter of the largest ball that can surely pass through a given structure without getting stuck in it. The results obtained are summarized in Table 2.

**Table 2:**
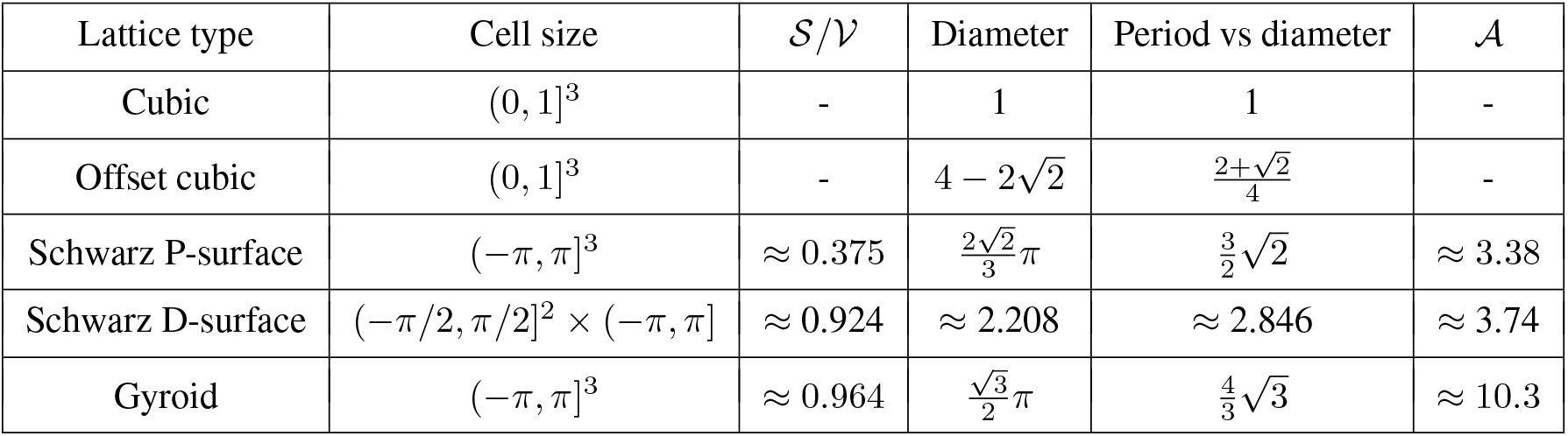
Results for the different structures considered, namely dimensionless periodicity used in the computation, ratio of surface area to volume of the periodic structure, diameter of the largest ball identified that will surely pass through the scaffold, ratio of the period of the scaffold to the diameter of the ball, ratio of the surface area in the reference cell vs that of the ball.

Table 2 also reports some information related to the surface area of the scaffolds that are useful for applications. In fact, as described in the Introduction, in order to have an efficient migration, cells need to have the possibility to attach to the substratum to be able to exert the necessary traction forces. For this reason, cell migration is favored in structures with a great ratio of surface area to volume, here denoted by 𝒮*/* 𝒱. Generally speaking, one can notice that triply periodic surfaces are characterized by high values of this ratio, particularly gyroids, which explains their relevance in the construction of biomimetic scaffolds. For uniformity, we take (−*π, π*]^3^ as the reference cell for the P-surface and the gyroid and 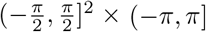 for the D-surface and calculate numerically the area of the surfaces in it. We find them to be 𝒮_*P*_ ≈ 93.1 for the Schwartz P-surface, 𝒮_*D*_ ≈ 57.3 for the Schwartz D-surface and 𝒮_*G*_ ≈ 239 for the gyroid, and therefore

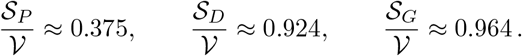

Actually, rather than using as reference a dimensionless reference length, in applications it is useful to refer to the size of the sphere that needs to enter the structure and report the ratio of the surface area of the structure with respect to the area of the sphere. This is given by the dimensionless number : 𝒜= 𝒮*/*(*πD*^2^) where *D* is the diameter of the biggest sphere that can pass through the structure. This leads to

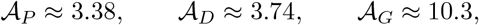

showing that gyroids have a much larger ratio compared to Schwarz surfaces.

Of course, in Table 2 we cannot report the same information for cubic lattices that in the idealized case are formed by lines.

## 3 Cubic Lattices

The case of cubic lattices represented in the first two rows of Table 1 is trivial. However, their discussion here is useful to explain the objectives pursued in this article and the methods used. In fact, it is clear that in the case of the standard cubic lattice, the largest ball that passes through the face of the reference cell is a ball with diameter *D* equal to the edge *l* of the cubic reference cell, see Figure 1a. When the sphere enters inside the cube, the largest sphere, touching the sides of the cubes, has a larger diameter. So, the bottleneck is the entrance into the cube, leading to a critical ratio between the diameter of the sphere and the size of the cube equal to 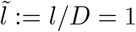.

**Figure 1:**
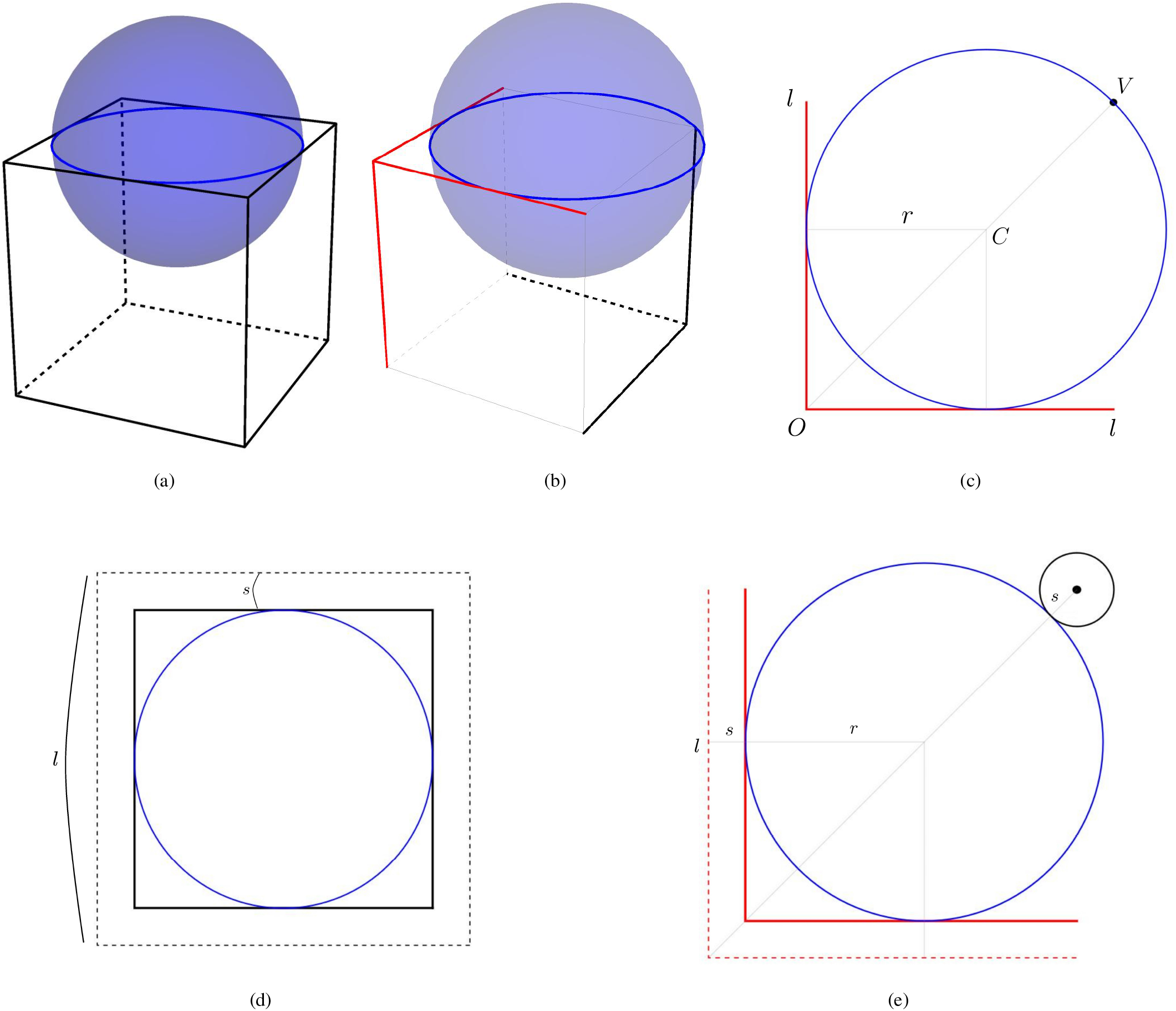
Sphere passing in a cubic lattice (a) and in a offset cubic lattice (b). Maximum circle passing through the face of the offset cubic lattice (c). In (d) and (e), corrections respectively for thickened cubic lattice and offset cubic lattice.

In the case of an offset cubic lattice represented in the second row of Table 1, the largest ball that can enter must have its equatorial circle touching two adjacent perpendicular edges of length *l* (in red in Fig. 1b) and the closest point *V* in that plane belonging to the dual lattice (in black in Fig. 1b). Due to symmetry, the center of the ball *C* lies on the diagonal joining *O* and *V* and

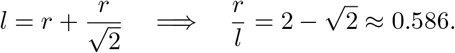

We remark that, trivially, the ball is bigger than the one obtained in the previous case.

Using the diameter of the sphere as reference, the dimension of the offset cubic lattice versus the diameter of the sphere must be such that

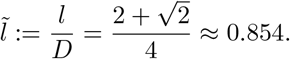

### Remark 3.1.

In the case of thickened cubic lattices, closer to realistic scaffolds, some obvious corrections need to be introduced in the above formulas. Regarding the standard cubic lattices, denoting by 2*s* the thickness of the edges of the lattice, i.e. every thickened edge is a cylinder of radius *s*, the largest ball passing through it will naturally have diameter *D* = *l* − 2*s* (see Figure 1d, that presents a face of the cube with the maximal circle of the sphere inscribed into the thickened face). Thus, 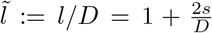. Regarding the offset cubic lattice, referring to Figure 1e, analogous computations and considerations lead to 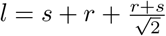, and therefore to

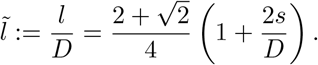

### 3.1 Triply-periodic minimal structures

The identification of the size of the largest sphere that will certainly pass through triply-periodic structures is based on the observation that when a sphere passes through the labyrinthine structure of a porous scaffold, it determines a tube generated by a central curve. The biggest tube that can pass through the lattice has a circular section that touches the surface in at least two points. However, as one can easily see from Figure 2, since we are working with triply periodic minimal surfaces with the symmetries mentioned in Section 2, configurations with an odd number of contact points (such as the first and the third in Figure 2) cannot guarantee symmetries with respect to two perpendicular axes or to permutations of the three coordinates. Therefore, the maximum circle cannot touch the surface in an odd number of points, and thus, as shown in the central picture in Figure 2, the contact points identify the diameter of the circle, and the normal at such points is directed toward another contact point with opposite normal.

**Figure 2:**
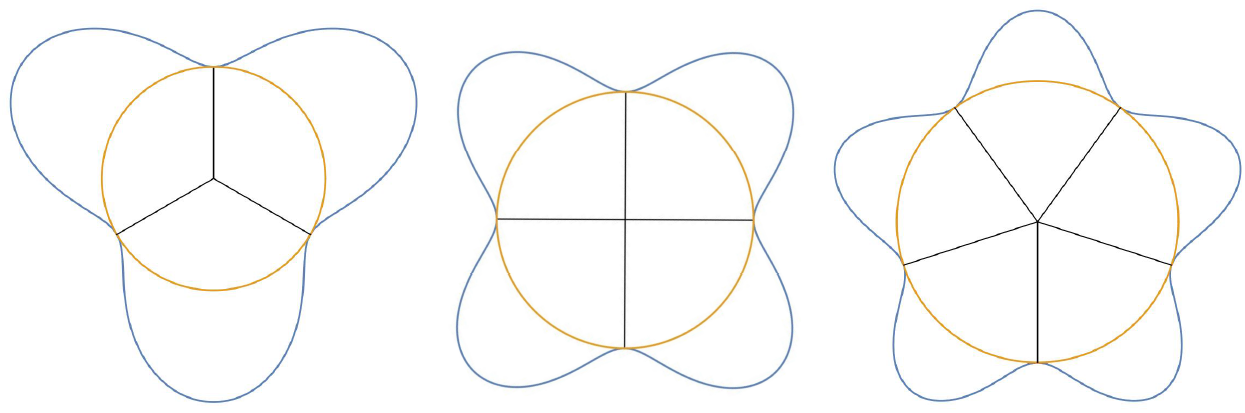
Circles inscribed in different holes determined by a different number of contact points (namely, 3,4,5). The first and the third configurations are symmetrical with respect to an axis (e.g., the vertical one), but not with respect to its orthogonal direction.

In this respect, we remark that in the case of the offset cubic lattice that is characterized by the presence of two nested structures in the periodic cell of side 2*l*, the reasoning above does not apply, although the reference cell (of side *l*) has a central symmetry. In fact, as shown in Figure 1e, the maximal circle has an odd number of contact points, two lying on the edges of the face of the reference cell and one is the point belonging to the edge perpendicular to the face, corresponding to three incident normal vectors. In fact, Figure 1e shows that the configuration of a face of this lattice has a symmetry with respect to the line *y* = *x*, but not with respect to the perpendicular direction.

So, our strategy is the following: for any pair of points *A* and *B* on the surface, we check whether the normal vector to the surface in *B* is directly opposite to the one in *A*. Among the possible pairs of points satisfying this criterion, we then choose the one with the minimal distance, such that the circle (great circle of the sphere) with diameter equal to this distance does not intersect the surface in other points.

In mathematical terms, denoted by Σ the surface given implicitly by *F* (**x**) = 0, we consider the set of pairs of points *A, B* ∈ Σ with coordinates **x**_*A*_ and **x**_*B*_ such that

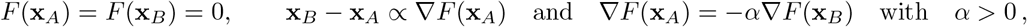

or equivalently

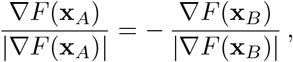

i.e. the two normal vectors to Σ in *A* and *B* are opposite and parallel to the line connecting the two points. In this set we then select the pair with minimum Euclidean distance between *A* and *B*. We then check that the maximum curvature of the surface in *A* and *B* is smaller than the curvature of the sphere, i.e. 2*/D*.

In this way we find the diameter of the largest sphere passing through that restriction of the porous scaffold. In particular, the fact that we look for the smallest passage leads to minimize the Euclidean distance between the two points, while looking for the largest ball implies that the endpoints of the diameter are precisely **x**_*A*_ and **x**_*B*_, i.e. that *A* and *B* belong to the surface and that the sphere is tangent to the surface with a curvature larger than that of the surface and therefore cannot be enlarged without violating non-interpenetration of the two bodies.

The steps we will then analyze are the following:

- Study of the suitable local expressions of *z* for the surfaces as functions of *x, y* in the reference cell, and of their domains;
- Computation of the normal vector to the surface;
- Identification of the set of point pairs that have opposite normals.
- Calculation of the Euclidean distance between such pairs of points and determination of the parameter that guarantees the minimal Euclidean distance.
- Verification that there is no interpenetration of matter between the structure and the sphere by computing the curvatures at the points of contact between the sphere and the surface: the maximum sectional curvature of the surface at those points must be less than that of the sphere.

#### Remark 3.2.

We remark that in this way the center of the ball will be the midpoint between the two points with opposite normals. Moreover, being the ball tangent to the surface, we will have that the diameter lies along the line that does not only join the two points, but on which also the two normals lie. This will be important in the following.

We advise, however, that this procedure might be sub-optimal in the sense that it assures that a sphere with diameter smaller than the one identified will certainly pass through. However, in geometries with interconnected surfaces and pores or with relevant size variability, it cannot exclude that there might be a path using larger pores that allows bigger spheres to pass. For example, in the case of the thickened offset cubic lattice, the procedure described above would identify the points on the open size of the reference cell, yielding a smaller *D* = *l* − 2*s* as in the cubic lattice with respect to the results discussed at the end of the previous section.

We underline that the maximal sectional curvature of a surface in a point is the maximum of the two principal curvatures *κ*_1_ and *κ*_2_. We recall that *κ*_1_ and *κ*_2_ can be easily obtained from the Gaussian curvature and the mean curvature that are the product and the mean of *κ*_1_ and *κ*_2_ and can be computed using standard formulas of differential geometry [5], through derivatives of the implicit function that defines the surface. We finally remark that the expressions used are approximations of proper Weierstrass representations of triply periodic minimal surfaces, so the mean curvature does not vanish in every point.

## 4 The Schwarz P-surface

The P-surface is described by the implicit relation

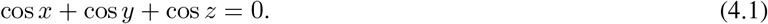

The reference cell is a cube of edge 2*π* that we will take as the set (−*π, π*]^3^. The origin is then a point of symmetry, as are the three axes and the three coordinate planes. Therefore, if 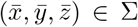 with 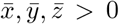, then there are 7 other points 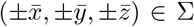 for all possible combinations of signs. In addition, any permutation will also belong to the surface, e.g. 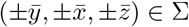.

### 4.1 Entering the Schwarz P-surface Neck

The sphere passing through the reference cell must clearly enter through one of the holes on the faces of the cube, having the surface an intuitive “interior” and “exterior” region. We choose the entrance with *z* = *π* and after observing that the normal vector to the surface at a point with *z*_*A*_ = *π* has a vanishing *z*-component, we look for the greatest circle in the plane *z* = *π* that can be inscribed, and therefore tangent, into the curve obtained by the intersection of the surface and the plane, i.e. cos *x* + cos *y* = 1. Due to symmetry, the center of the circle will be the point **x**_0_ = (0, 0, *π*). We can then reduce the study to the first quarter of the plane *xy*, i.e *x, y* ∈ [0, *π/*2], as shown in Fig. 3c.

**Figure 3:**
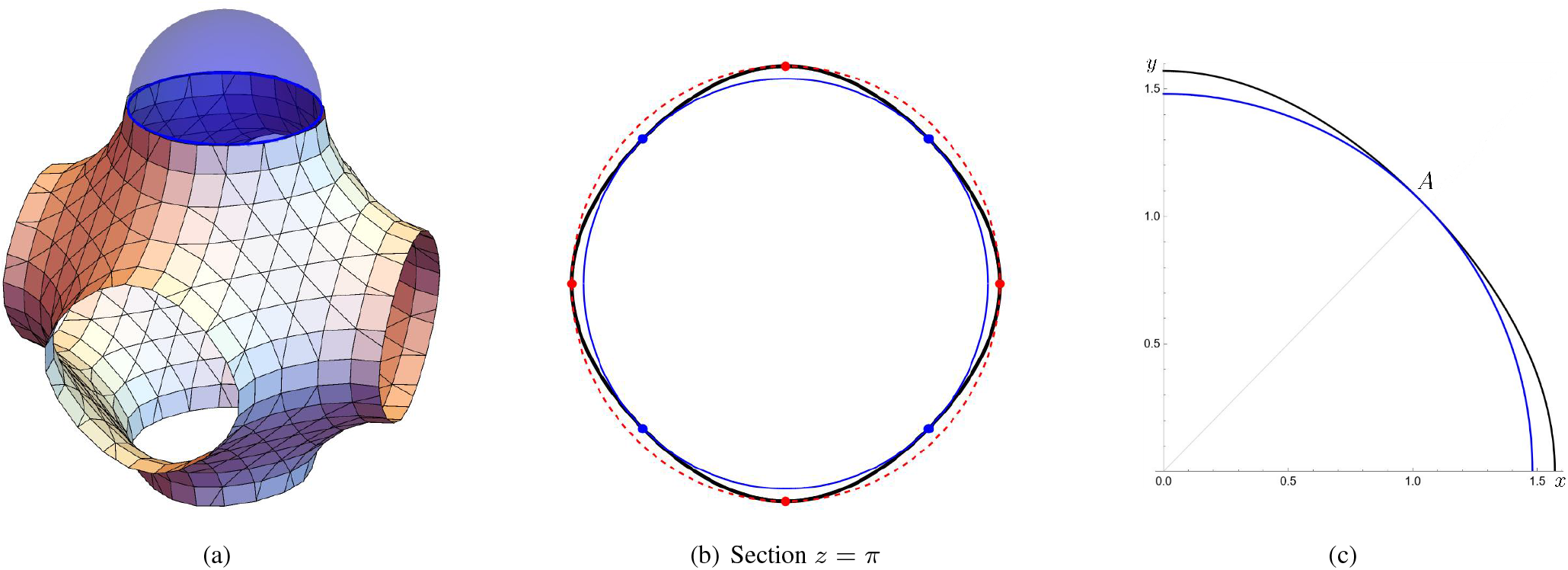
(a) Sphere entering the P-surface structure from the neck in *z* = *π*. (b) Intersection of the P-surface with the plane *z* = *π* in black and circle with 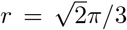 in blue (which is admissible) and with *r* = *π/*2 in dashed red (which is not admissible) and the corresponding contact points. (c) Notation used in Section 4.1.

The fact that the normal vectors to the two curves must be colinear implies that

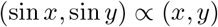

and thus if *x, y ≠* 0,

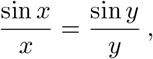

that in 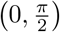 is only solved by *x* = *y* due to the monotonicity of 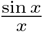 in that interval. Hence, the contact point *A* is given by 2 cos *x* = 1, i.e. 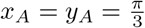. The radius of the circle is then 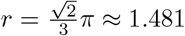. In order to be sure that locally the circle is inscribed in the curve, we notice that the curvature of

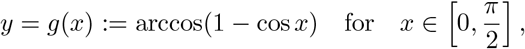

in 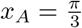 is

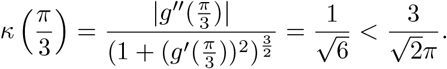

Checking the maximal sectional curvature in the contact point 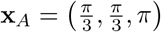, we have that

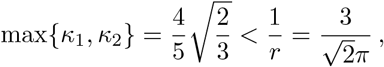

and thus the result is physically admissible. The other contact point will read 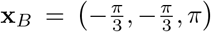 and we obtain 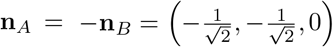, see blue points in Figure 5a.

Trivially, the points with vanishing *x* or *y* that were not considered above have a distance from the origin equal to *π/*2, greater than that of the point *A*, just determined (see Figure 3b and red points in Figure 5a). In addition, the curvature of *g*(*x*) in such points (with vanishing *x* or *y*) is larger than that of the sphere, since 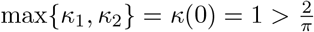 and therefore the sphere would intersect the porous structure. So, these points are anyway not acceptable.

We remark that the point of tangency *A* is precisely the nearest point of *g*(*x*) to the origin, that is,

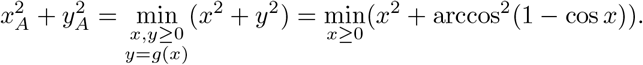

In conclusion, the diameter of the largest sphere that can enter the structure is

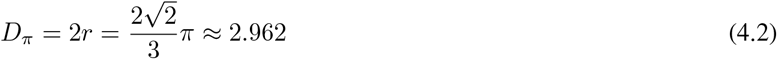

and the ratio of the periodicity length to the diameter is

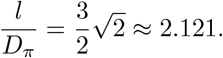

### 4.2 Inside the Schwarz P-surface

We expect that once entered, the ball will find a less constricting region. In order to prove this, let us first examine the case in which *z*_*A*_ = 0. As for the case *z*_*A*_ = *π* treated in the previous subsection, the normals to the surface at points with *z*_*A*_ = 0 have a vanishing third component and therefore also *z*_0_ = 0.

Proceeding in the same way as in the previous subsection, we have to find the circle inscribed in the curve cos *x* + cos *y* + 1 = 0. One can prove that the points with opposite normals have again *x* = *y*, but this time 2 cos *x* = −1 and, therefore, *x* = *y* = 2*π/*3, as shown in Figure 4a. The two contact points are 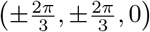 and the normal vectors read 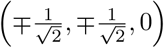, see Figure 5b.

**Figure 4:**
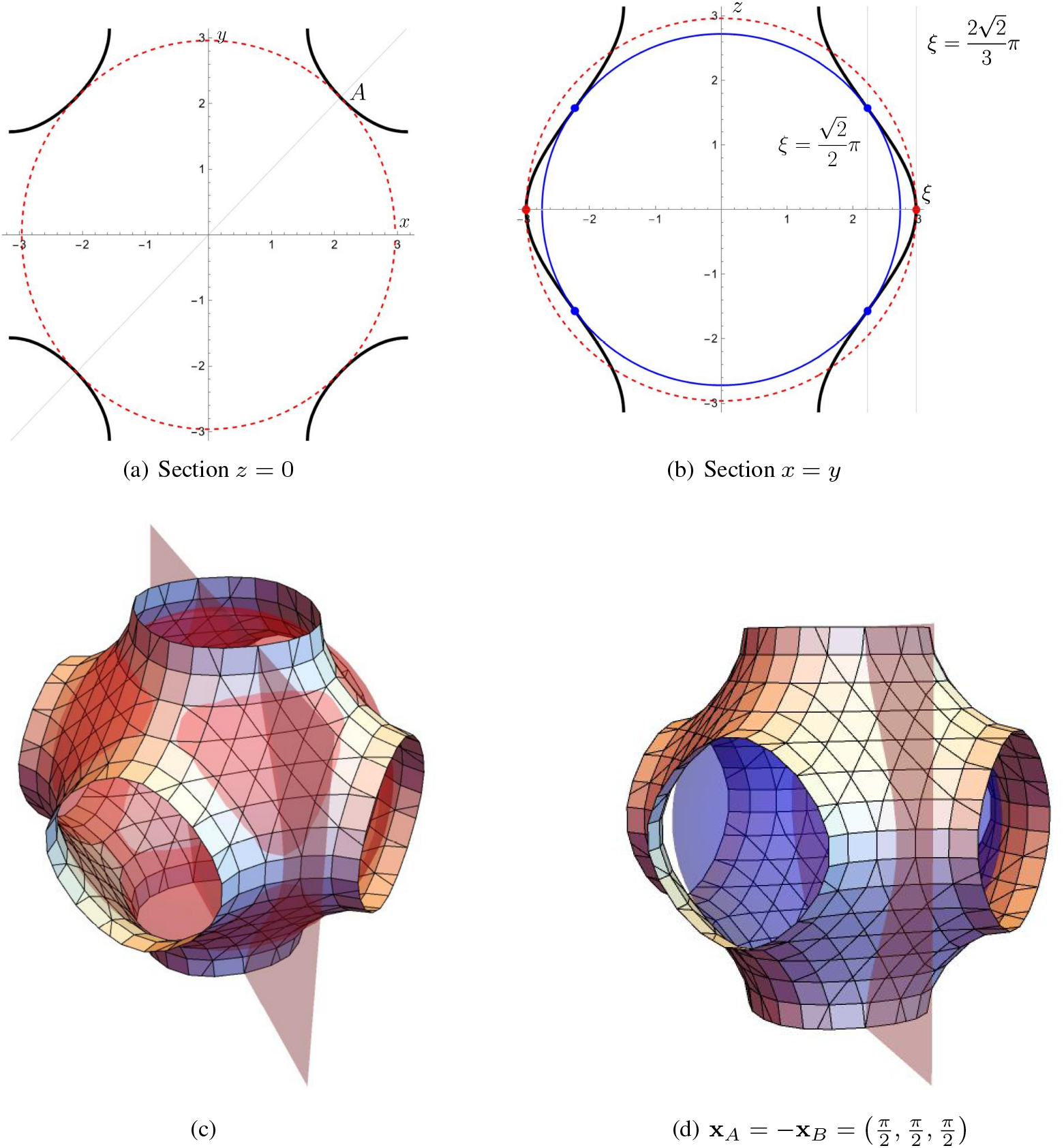
Sections of the Schwarz P-surface with *z* = 0 in (a) and with the plane 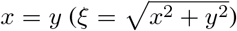 in (b). The surface is in black and the non intersecting circles in blue while the intersecting ones are in dashed red. (c) Interpenetration of matter between the non admissible sphere and the surface when the contact point is 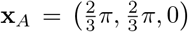, as put in evidence by the shaded red area. (d) The sphere and the surface with 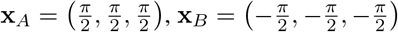.

**Figure 5:**
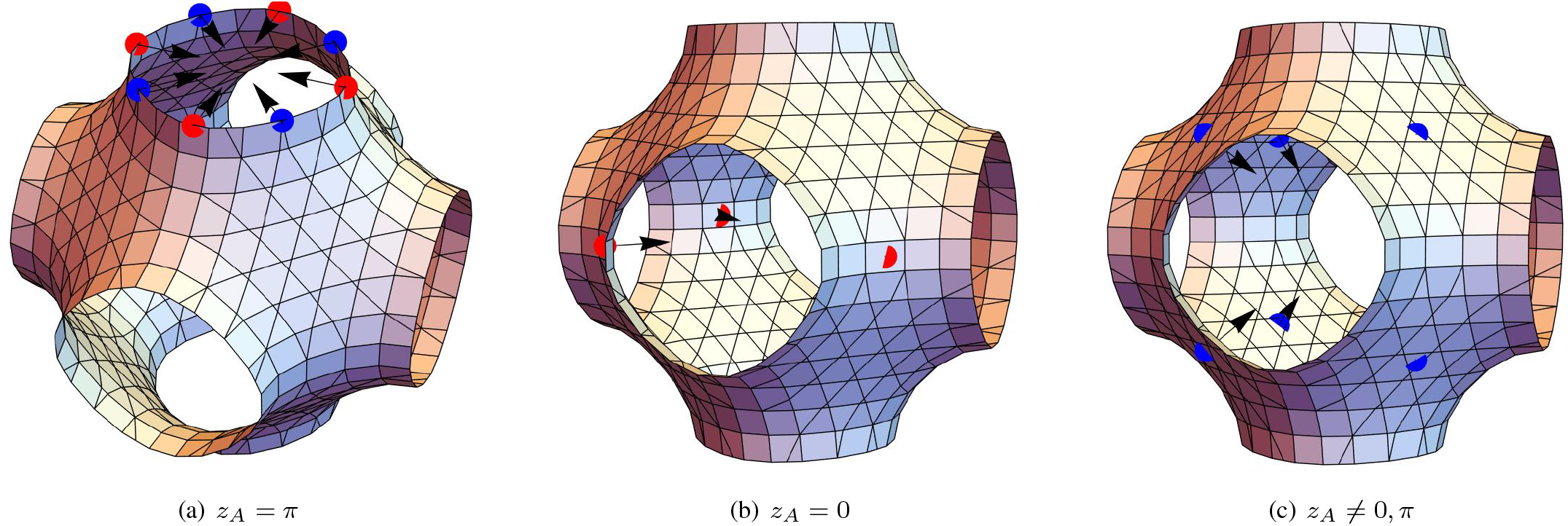
Contact points belonging to admissible and non-admissible spheres, respectively in blue and red. (a) At the entrance *z* = *π*, the contact points 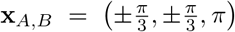 and the symmetric pair 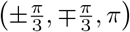 are admissible, while 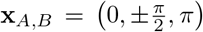 and the symmetric pair 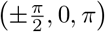 are not; (b) for 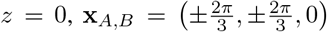 and the symmetric pair 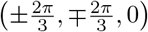 are all not admissible; (c) 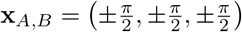 and all the symmetric pairs obtaining changing one or more signs are admissible, but have a distance larger than in (a). For all the points, also the normal vectors are displayed, directly opposite for every pair.

We have 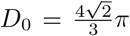, which is bigger (the double) than the one already found for *z* = *π*. This means that such a ball would not pass through the hole in the faces of the Schwarz P-surface (e.g., at *z* = ± *π*) to get to the center of the reference cell. Though, of course, we have all cues to exclude this case, we observe, in addition, that in the section *x* = *y* (drawn in Figure 4b) in the contact point 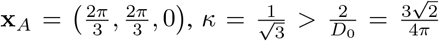, that implies that in the plane *x* = *y* the tangent sphere will intersect the structure, as shown by the shaded area in Figure 4c, which is not allowed. Indeed, looking at the sectional curvatures in the contact point 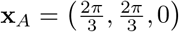, one has

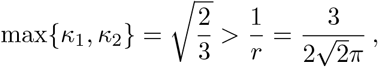

which implies that there is interpenetration of matter. Due to symmetry, the same holds for the planes *x* = 0 and *y* = 0. So, in conclusion, there is no admissible tangent sphere with contact points that have either *x*_*A*_ = 0, or *y*_*A*_ = 0, or *z*_*A*_ = 0.

It remains to look at the case with *z*_*A*_ ≠ 0, *π*. By symmetry and without loss of generality we can assume that one of the two points of contact has *x*_*A*_, *y*_*A*_, *z*_*A*_ *>* 0 with normal

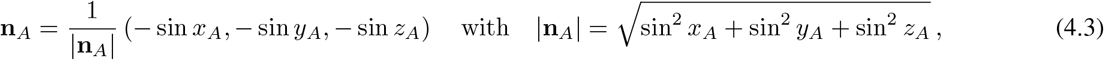

so that **n**_*A*_ has three negative components. The points on the surface with all positive components are those in the opposite octant, i.e. *x*_*B*_, *y*_*B*_, *z*_*B*_ *<* 0.

It is clear by inspection that the point *B* ∈ Σ with **x**_*B*_ = − **x**_*A*_ has opposite normal vector. Later on, we will prove that there are no other candidate points. For the moment let us focus on the analysis of this case which also implies that **x**_0_ = **0**.

#### 4.2.1 Opposite Contact Points

Expliciting (4.1) for *z >* 0 gives

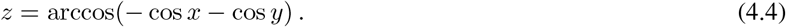

Hence, the two opposite points are

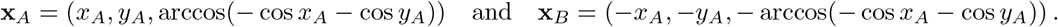

Their distance (squared) is given by

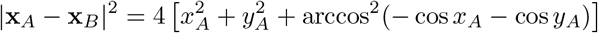

that needs to be minimized for *x*_*A*_, *y*_*A*_ *>* 0. This leads to

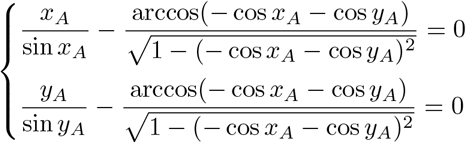

which implies that

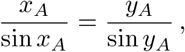

that again in (0, *π*) is only solved by *x*_*A*_ = *y*_*A*_.

Therefore,

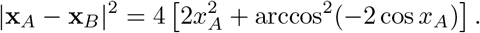

By a standard study it can be checked that the distance is minimized when *x*_*A*_ = *π/*2. In conclusion,

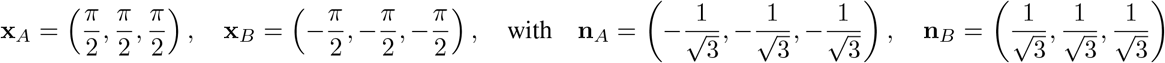

and

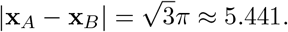

So, though in this case the sphere would not cross the surface as before (both principal curvatures vanish), the distance between the two points (drawn in Figure 5c) is larger than the distance 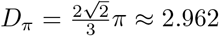 found in (4.2) between the blue opposite points in Figure 5a. So, also this sphere would not pass through the neck, say, in *z* = *π*.

#### 4.2.2 Other Possible Contact Points

Recalling (4.4), the upper fold of the surface with 0 *< z < π* is given by *z* = arccos(− cos *x* − cos *y*) and the lower one with −*π < z <* 0 by *z* = − arccos(− cos *x* − cos *y*). The two functions have the same domain of existence

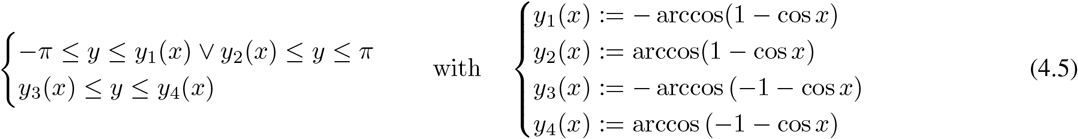

with *x, y* ∈ (− *π, π*] drawn in gray in Figure 6.

**Figure 6:**
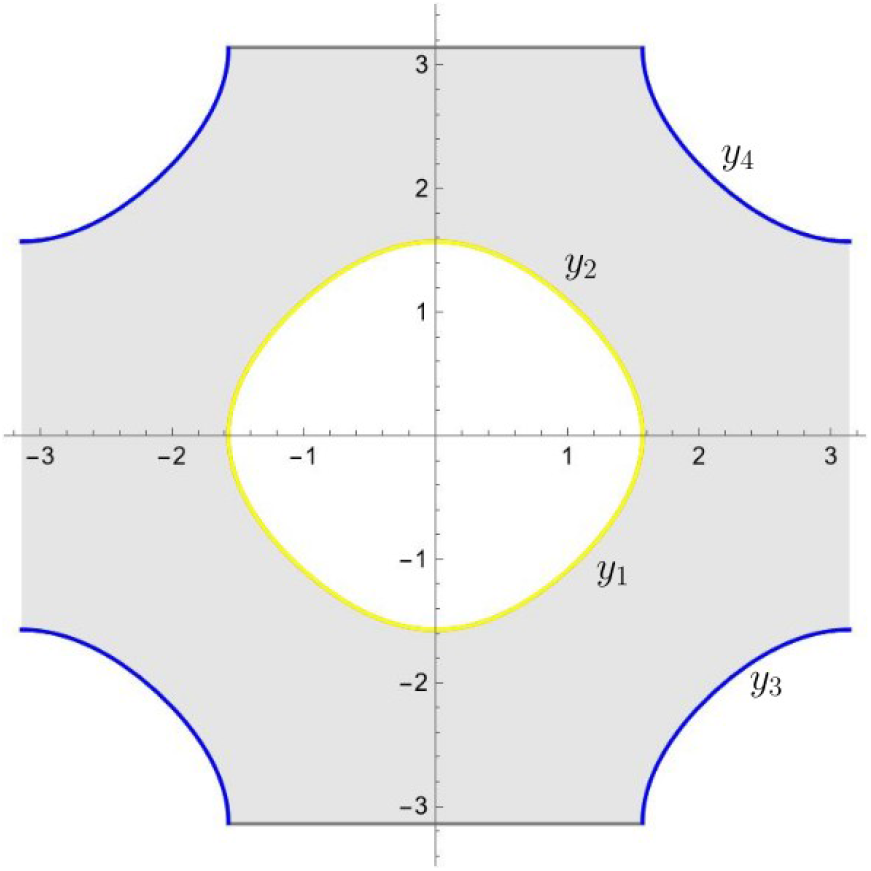
The two conditions in (4.5) identify the regions outside the yellow curve (identified by the top equalities) and inside the blue line (identified by the bottom equalities). The domain of existence of the function is in grey. The functions *y*_1_(*x*), *y*_2_(*x*), *y*_3_(*x*), *y*_4_(*x*) are also used in Appendix A.

Eliminating *z* from (4.3) gives

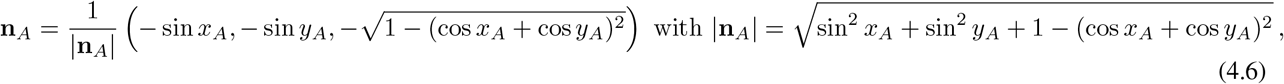

and similarly

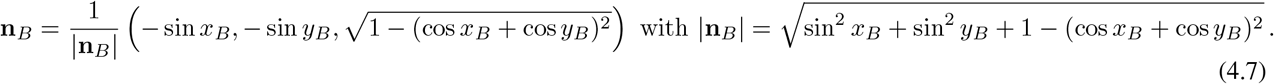

There are now several cases that must be studied. However, many of them can be excluded by symmetry and the analysis made in Subsections 4.1 and 4.2. Their complete analysis, that follows the strategy mentioned in Subsection 3.1, is carried out in the Appendix A. We thus only report here the analytical proof when the two points belong to opposite octants, e.g. *x*_*A*_, *y*_*A*_ *>* 0 and *x*_*B*_, *y*_*B*_ *<* 0, showing that there are no other points that must be taken into account in addition to those described in Subsection 4.2.1.

The request that the three components of the normal vectors must be opposite leads to the system

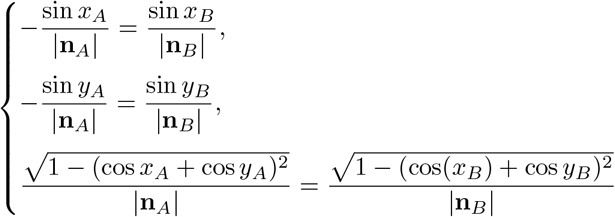

or

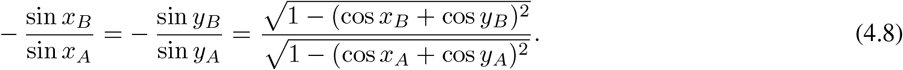

The first relation of (4.8) tells us that

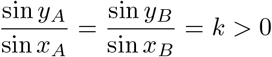

and thus *A* and *B* can be such that:

1. *y*_*A*_ = arcsin(*k* sin *x*_*A*_) while *y*_*B*_ = arcsin(*k* sin *x*_*B*_) (or analogously *y*_*A*_ = *π* − arcsin(*k* sin *x*_*A*_) and *y*_*B*_ = − *π* − arcsin(*k* sin *x*_*B*_), where this last expression comes from the fact that *k >* 0 and *x*_*B*_ *<* 0); since by assumption *x*_*A*_ *>* 0 and *x*_*B*_ *<* 0, the relation between the first and the third term of (4.8) has both the left and right terms positive, so we can square both to obtain

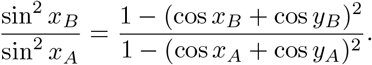 This leads to

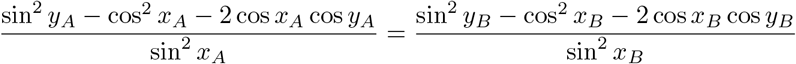

and substituting *y*_*A*_ and *y*_*B*_

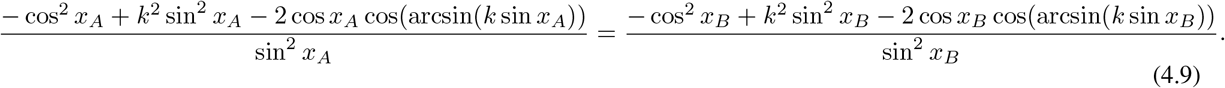 Thus, denoting

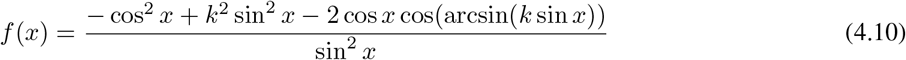

the preceding equality is analogous to finding two abscissas for which the even function *f* has the same value. The function *f* (*x*) is even, so we can restrict the study to the interval *x* ∈ (0, *π*) or, if *k >* 1, to its subintervals where *k* sin *x* ∈ [−1, 1], i.e.,

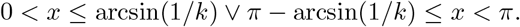 In this interval *f* is monotonically increasing (and therefore monotonically decreasing for *x <* 0). In fact,

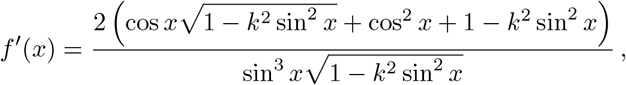

which is naturally positive because we are working in the domain with 1 − *k*^2^ sin^2^ *x >* 0 and *x* ∈ (0, *π*). This implies that (4.9) is only satisfied by *x*_*B*_ = −*x*_*A*_ and then *y*_*B*_ = −*y*_*A*_, which falls in the case studied in Section 4.2.1.
2. *y*_*A*_ = arcsin(*k* sin *x*_*A*_) and *y*_*B*_ = − *π* − arcsin(*k* sin *x*_*B*_) (or analogously *y*_*A*_ = *π* − arcsin(*k* sin *x*_*A*_) and *y*_*B*_ = arcsin(*k* sin *x*_*B*_)); we can proceed as before, looking for abscissas such that *f* (*x*_*A*_) = *f* (*x*_*B*_) where *x*_*A*_ *>* 0 and *x*_*B*_ *<* 0. But *f* (*x*_*A*_) and *f* (*x*_*B*_) are precisely the two branches of the preceding *f*, restricted respectively to *x* ∈ [0, *π*] and *x* ∈ [−*π*, 0] and then we fall again in the preceding case.

Thus, we could not find any couple of points with opposite normal vectors and satisfying Remark 3.2 that have distance less than *D*_*π*_ in (4.2). In conclusion, the largest sphere that can pass through the Schwarz P-surface with periodicity 2*π* has diameter 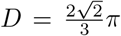, or viceversa if we want to be sure that a ball of diameter *D* passes through a Schwarz P-surface its period *l* must be such that

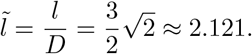

## 5 The Schwarz D-surface

The Schwarz D-surface is described by the implicit relation

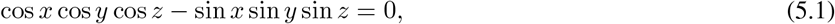

and the unit normal **n** in a generic point is

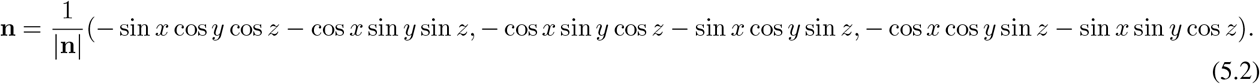

The symmetries of the surface allow us to restrict the study to the reference cell 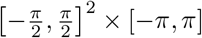.

We will first consider in Section 5.1 what happens when either *x* or *y* belong to the boundary of this reference cell, specifically when 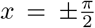 or 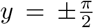. In this case, Eq. (5.1) trivially becomes sin *y* sin *z* = 0 in the former case, or sin *x* sin *z* = 0 in the latter. So, in the former case the points belonging to the Schwarz D-surface are such that *z* = 0, ± *π* for any *y*, or *y* = 0 for any *z*, and similarly (replacing *y* with *x*) for the latter case.

Away from the boundary of the (*x, y*)-domain, i.e. for 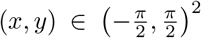, we find it more convenient to introduce 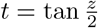 rather than expliciting (5.1) as tan 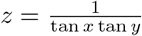 In this case, we can rewrite (5.1) as

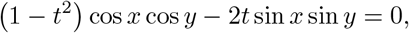

that is solved by

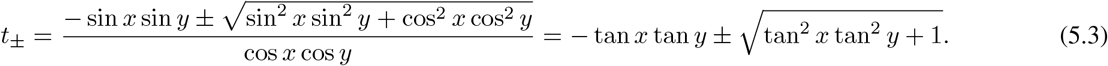

Thus, introducing 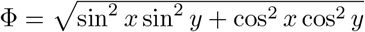, the two folds of the surface will be given by

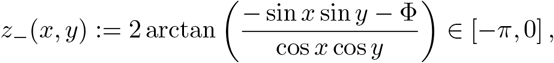

and

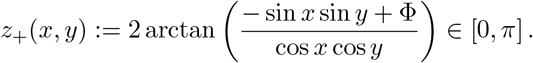

This relation will be the basis of the discussion in Section 5.2.

We also underline that the starting implicit function defining the surface is regular, since it is a polynomial function of sines and cosines. Then, the normal vector **n** is well-defined and continuous everywhere, also on points with cos *x* cos *y* = 0, where the components of the normal vector are defined as continuous extensions on the two folds.

### 5.1 Contact points with *x* or *y* on the border of the reference cell

In this case either cos *x* or cos *y* vanish nullifying the denominator of (5.3). This condition identifies for instance the four points

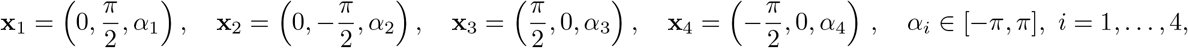

with normal

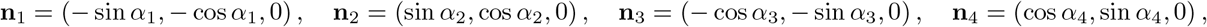

respectively. However, since *n*_*z*_ = 0 and we are looking for directly opposite normals, the two points must lie on the same horizontal plane, leading to the same value of *α*. We remark that, for example, **x**_3_ is obtained by **x**_1_ switching the *x* and *y* axes. The same holds for any switch of the axes. Therefore, because of this symmetry, it will be enough to restrict to this case.

Clearly, with *α*_1_ = *α*_2_, **n**_1_ = − **n**_2_ and the normals are directly opposite and aligned to **x**_2_ − **x**_1_ when *α*_1_ = *α*_2_ = 0, ±*π*. Similarly, for **x**_3_ and **x**_4_. In these cases the distance between the two points is *D*_1_ = *π* (we do not plot an example of this case).

On the other hand, the distance between the other possible couples, e.g. **x**_1_ and **x**_3_ with *α*_1_ = *α*_3_, is 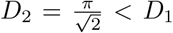. In this case (and by symmetry in the others) the normal vectors are opposite just when sin *z* = − cos *z*, i.e. 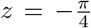, so the two candidate points with distance *D*_2_ will be 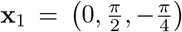 and 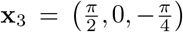 (see Figure 7a) with normals 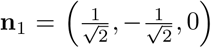 and 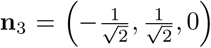. However, checking the maximal sectional curvature in both points, we have 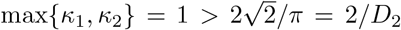, so interpenetration of matter locally occurs and the couple of contact points is not admissible, as shown in Figure 7b.

**Figure 7:**
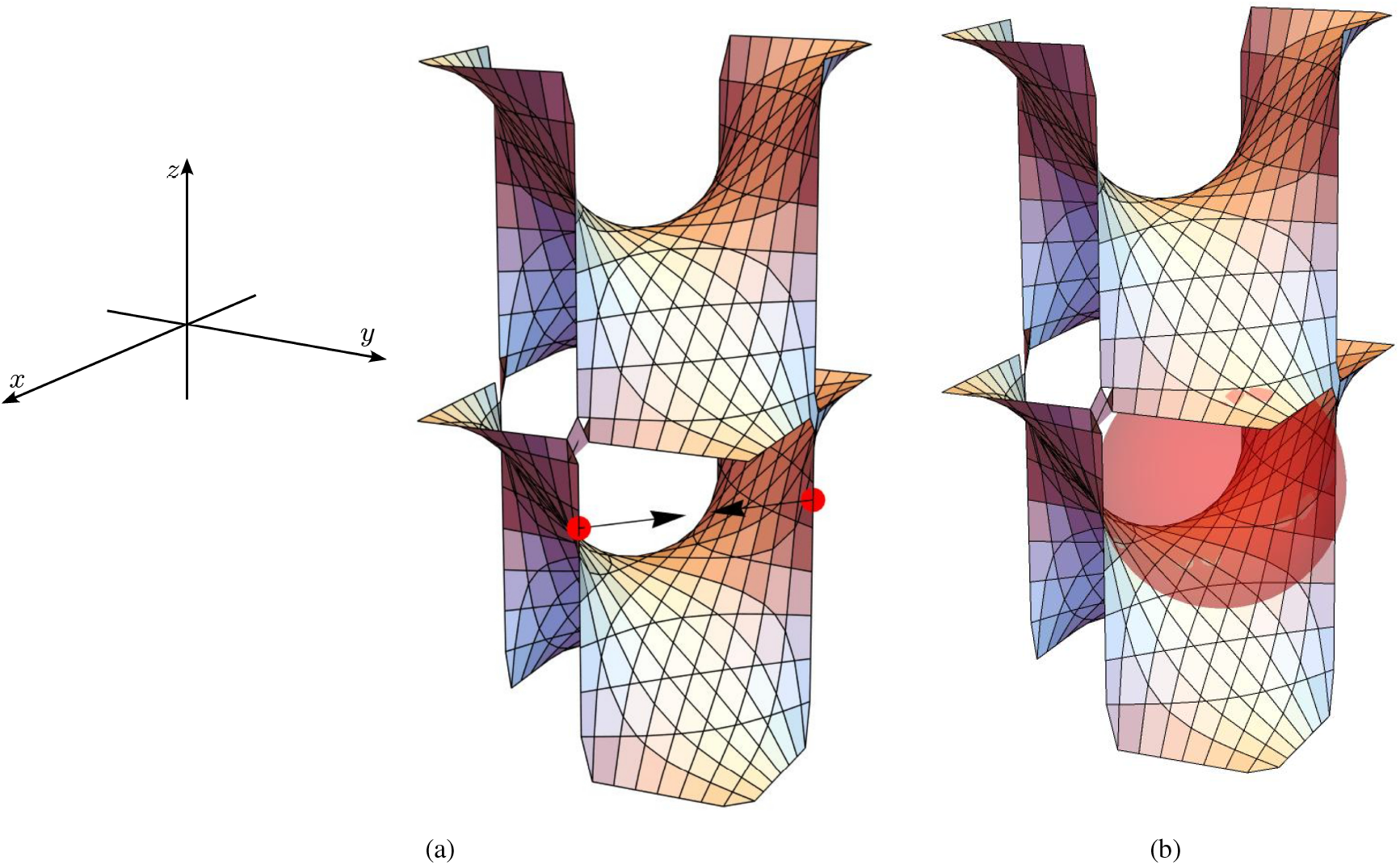
(a) Couple of points 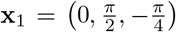 and 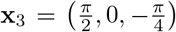 with opposite normals 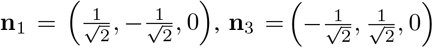 with minimal distance when either 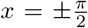 or 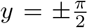. (b) Interpenetration of matter between the surface and the candidate sphere, as shown by the little shaded red area in the upper fold

### 5.2 Contact points on different folds

Having considered in the previous section the case in which either *x* or *y* are equal to *π/*2, we will here consider the case cos *x* cos *y*≠ 0 and therefore Φ *>* 0 and Eq. (5.3) well defined. In this case, the third component of the normal vector for a point on the lower fold is

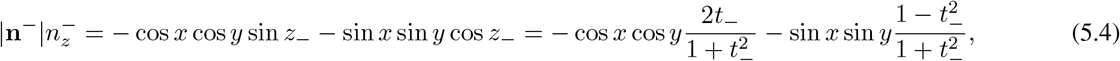

that with standard computations can be simplified to 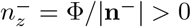. Similarly,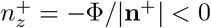.

This means that the two points cannot belong to the same fold and then we are going to look for the two points *A* and *B* belonging respectively to the fold *z*_−_ and *z*_+_, i.e. *z*_*A*_ = *z*_−_(*x*_*A*_, *y*_*A*_) and *z*_*B*_ = *z*_+_(*x*_*B*_, *y*_*B*_).

Proceeding as in (5.4) gives

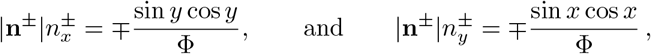

and therefore

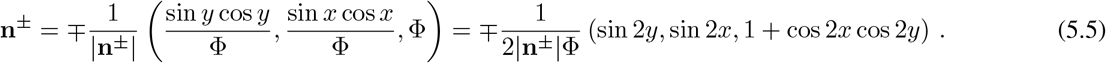

The study of the sign of the components tells that the two contact points must have (*x*_*A*_, *y*_*A*_) and (*x*_*B*_, *y*_*B*_) that belong to the same quadrant, e.g. the first without losing in generality, so *x*_*A*_, *y*_*A*_, *x*_*B*_, *y*_*B*_ ≥ 0. Hence, the two normals at the points *A* and *B* are opposite when

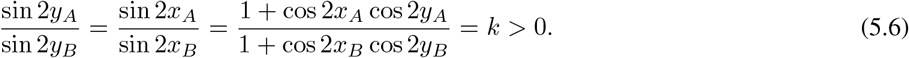

Before looking for the solution of the equation above, we observe that if, say, *x* = 0 for a point on the surface, then either 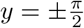 or 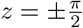. So, we fall back to the case already discussed (and excluded) in the previous section.

We will now prove that the set of equations in (5.6) has solutions only if *k* = 1, and

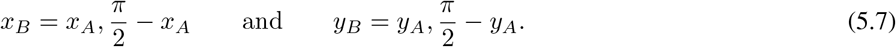

In fact, if for example we take *k <* 1 and work in 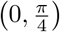, then the solution of the first equation would be such that *x*_*A*_ *< x*_*B*_ and *y*_*A*_ *< y*_*B*_. But then cos 2*x*_*A*_ *>* cos 2*x*_*B*_ and cos 2*y*_*A*_ *>* cos 2*y*_*B*_ and the last ratio is greater than 1 leading to an absurd. The other cases work similarly, and so *k* = 1 leading to the solutions in Eq. (5.7).

If *x*_*A*_ = *x*_*B*_ then one has that the normal must have *n*_*x*_ = 0, leading to *x*_*A*_ = *x*_*B*_ = 0 or 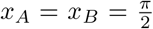 that have been already excluded. Similarly, if *y*_*A*_ = *y*_*B*_, then *n*_*y*_ = 0 and therefore *y*_*A*_ = *y*_*B*_ = 0 or 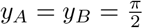.

It remains to consider the case 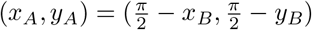, so that

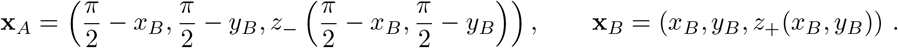

The requirement that **x**_*B*_ − **x**_*A*_ and **n** must be collinear gives

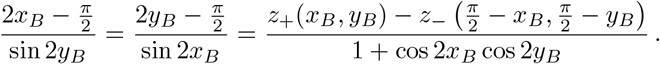

The first equality leads to 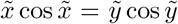 where 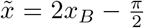 and 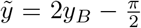, which in 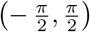 is only satisfied for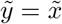, i.e. *y*_*B*_ = *x*_*B*_, because 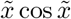 is a strictly increasing function there.

The second equality cannot be easily solved, so we compute the distance between *A* and *B*, minimize it, and then check if our solution satisfies the second equality for collinearity. The squared distance rewrites as

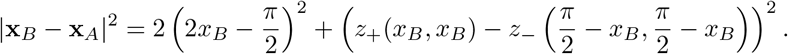

This function depends now on one variable only and thus we can minimize it, finding numerically 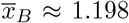. Moreover, the equality

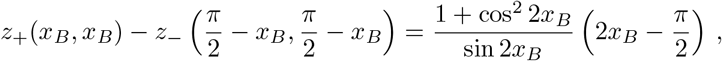

i.e. the second equality for the collinearity is satisfied by 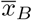.

Then, **x**_*A*_ = (0.373, 0.373, − 1.722) and **x**_*B*_ ≈ (1.198, 1.198, 0.152), and their distance is *D*_3_ ≈ 2.208 *< D*_2_ that is slightly smaller than *D*_2_ ≈ 2.221. The normal vectors are **n**_*A*_ ≈ (0.374, 0.374, 0.849) and **n**_*B*_ ≈ (− 0.374, −0.374, − 0.849), see Figure 8a. Moreover, the normal vectors are aligned with the segment joining *A* and *B* and indeed **x**_*A*_ + *D*_3_**n**_*A*_ = **x**_*B*_.

**Figure 8:**
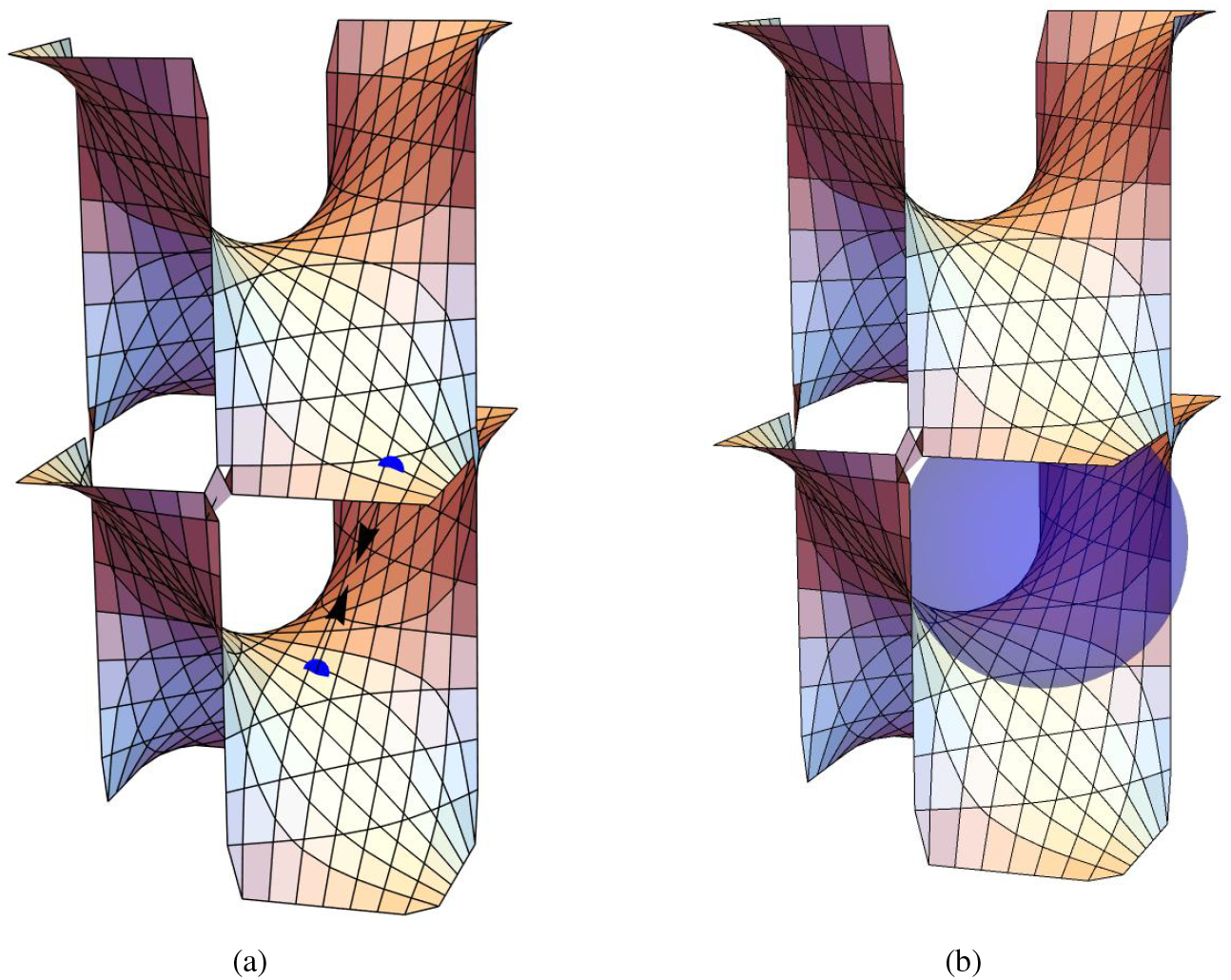
(a) The points *A* and *B* with opposite normals and belonging to different folds. (b) The ball having diameter *D*_3_ ≈ 2.208.

We also check the maximal sectional curvature, that is max {*κ*_1_, *κ*_2_ } ≈ 0.81 *<* 2*/D*_3_, so locally there is no interpenetration of matter and the ball is admissible, see Figure 8b.

Using the diameter of the sphere as reference, the dimension of the periodic cell must be

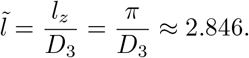

## 6 The gyroid

The surface is described by the implicit relation

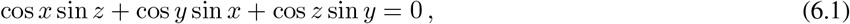

that, in view of expliciting it with respect to *z*, can be rewritten as *a* sin *z* + *b* cos *z* + *c* = 0 with *a* = cos *x, b* = sin *y*, and *c* = sin *x* cos *y*. As already mentioned, the reference cell is a cube of edge 2*π* that we will take as the set (− *π, π*]^3^.

We will first look at the case in which *b* = *c*, that is, when tan *y* = sin *x*, corresponding to the black curves in Figure 9. In this particular case, the points on the gyroid are such that

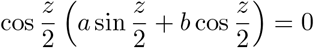

which gives *z* = *π* or (if *a*≠ 0) 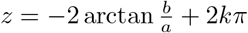. Since *z* ∈ (−*π, π*], then *k* = 0 and we can write the latter as

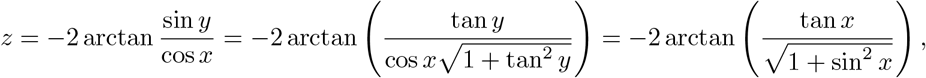

where we used again the fact that tan *y* = sin *x*.

**Figure 9:**
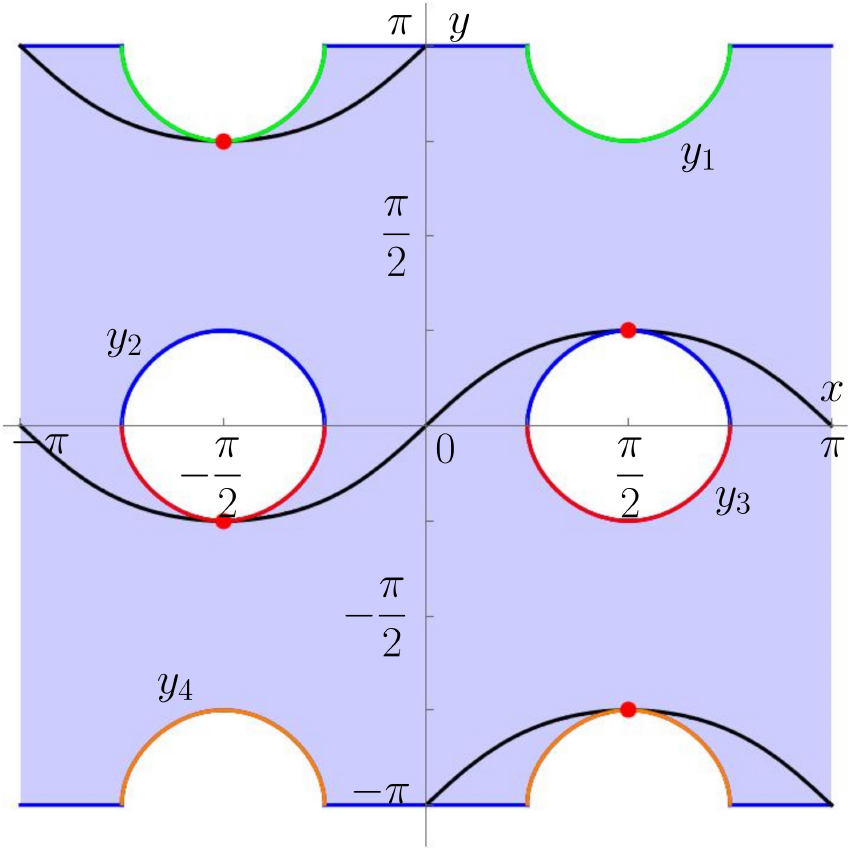
The domain of existence of the two folds of the gyroid written as *z* = *z*_*±*_(*x, y*). The boundary is determined partially by the lines *y* = ± *π* and by the four curves *y* = *y*_1_(*x*) = *π* − *β*(*x*) in green, *y* = *y*_2_(*x*) = *β*(*x*) in blue, *y* = *y*_3_(*x*) = − *β*(*x*) in red, *y* = *y*_4_(*x*) = − *π* + *β*(*x*) in orange, where *β*(*x*) is defined in Eq. (6.5). The black curves represent the points such that tan *y* = sin *x*, the red dots the points belonging to both the boundary of the domain of the folds and the black curves.

When *c*≠ *b* it is convenient to introduce 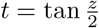 and rewrite (6.1) as

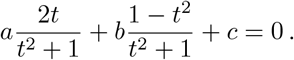

This equation is solved by

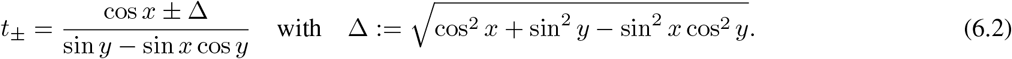

where the denominator does not vanish because we are dealing with the case *c*≠ *b*.

Then, there exist two possible expressions for *z* ∈ (−*π, π*] that are

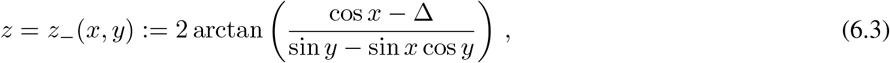

and

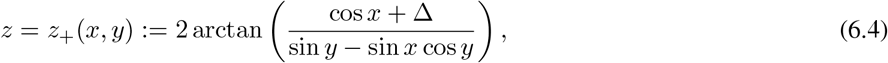

that will be respectively called the lower and upper folds.

We now have to look carefully at the domain of the expressions *z*_*±*_ in (*x, y*) ∈ (− *π, π*]^2^. The positivity of the radicand requires

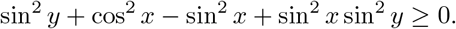

If 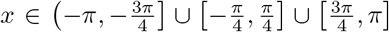, then | cos *x*| ≥ | sin *x*| and the inequality is always satisfied. Otherwise, we must limit ourselves to

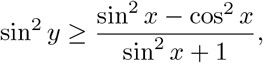

that implies

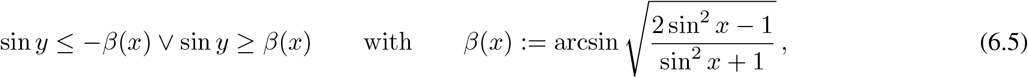

and thus when 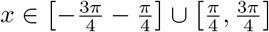 it must be *y* ∈ [*y*_4_(*x*), *y*_3_(*x*)] ∪ [*y*_2_(*x*), *y*_1_(*x*)] where

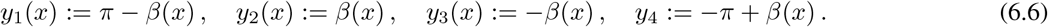

The domain of the folds is represented in blue in Figure 9.

Before starting the discussion on the normals, we remark that the starting implicit function *F* defining the gyroid is regular, since it is a polynomial function of sines and cosines. Then, we can compute its gradient, that is continuous. Moreover, a simple study shows that ∇*F*≠ 0 on the gyroid. Thus, the normal vector **n** to the surface is well-defined and continuous everywhere, also on points with sin *y* = cos *y* sin *x*, where the components of the normal vector are defined as continuous extensions on the two folds.

The unit normal **n** to the surface in a generic point of the gyroid is

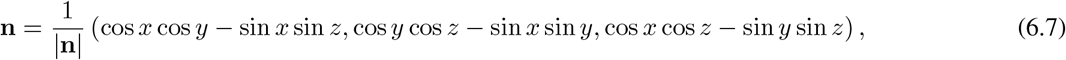

with 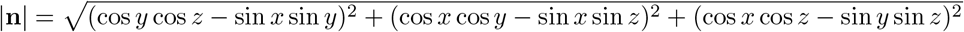.

Evaluating the third component on the upper fold gives

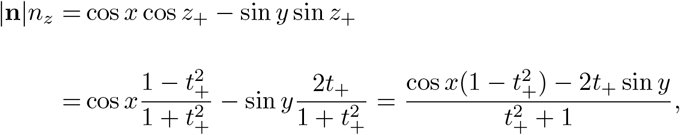

that by substituting *t*_+_ from (6.2), with tedious but standard simplifications, leads to *n*_*z*_ = − Δ*/* |**n**| . Thus, the *z*-component of the normal is obviously negative, since Δ is a square root. Similarly, on the lower fold *n*_*z*_ = Δ*/* |**n**|, see Figure 10. Hence, at the boundary between the lower and upper folds, where Δ = 0, the normal must have *n*_*z*_ = 0, as can be expected because there one has a double root and the surface bends over, which further supports the distinction between the lower and upper fold.

**Figure 10:**
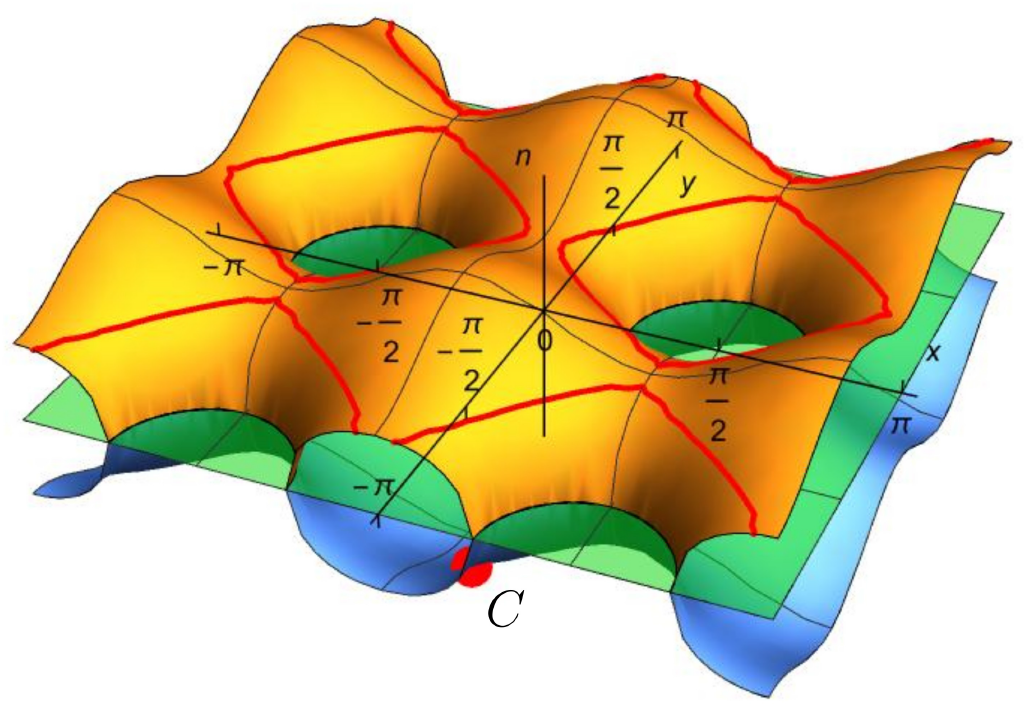
The symmetric functions 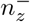 in orange and 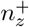 in blue. In green, the plane *n*_*z*_ = 0. In red, an example of level set 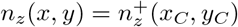 where *C* is the red point with coordinates 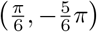.

### 6.1 Points with opposite normal vectors: Two points on the same fold

The only possibility for having two points with opposite *z*-components on the same fold is that *n*_*z*_ = 0 for both points *A* and *B*. This implies that the two points must belong to the same plane *z* = const with a vanishing Δ, i.e. cos^2^ *x* + sin^2^ *y* − sin^2^ *x* cos^2^ *y* = 0. Therefore, *z*_+_ = *z*_−_ := *z*_0_ and the projections of the two candidate points *A* and *B* on the plane *z* = 0 have to lie on the inner boundary of the domain in Figure 9, i.e. on the curves *y*_1_(*x*), *y*_2_(*x*), *y*_3_(*x*), *y*_4_(*x*), with 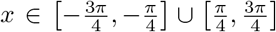, that were exactly identified by the limit condition Δ = 0. The fact that Δ = 0 also allows to simplify

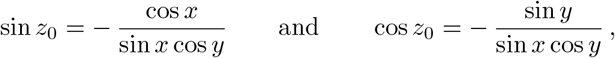

so that

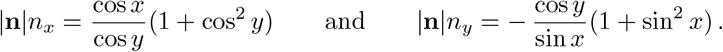

We have now different possibilities: the coordinates of the points can satisfy the same function (i.e., *y*_*A,B*_ = *y*_*i*_(*x*_*A,B*_)) or two different ones. Let us analyze the different cases, stressing again that the two points have the same *z*-coordinate.

#### 6.1.1 Projections on the Same *y*-function

Considering the case in which the two projections of *A* and *B* on the plane *z* = 0 lie on the same *y*-function, by symmetry, without losing in generality, we can assume that they both lie on *y* = *y*_2_(*x*) = *β*(*x*). Then for both of them

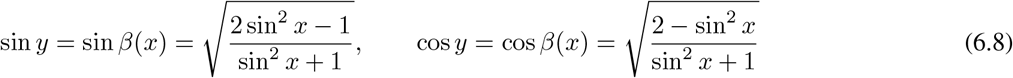

and

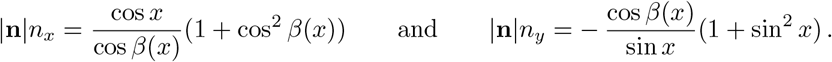

Moreover,

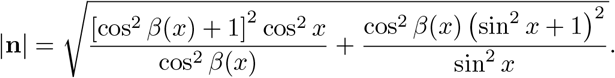

Clearly, *β*(*x*) is even, and since both sin *β*(*x*) and cos *β*(*x*) are even functions and, restricted to the interval 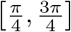, are symmetric with respect to the line 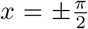, then *n*_*x*_ is even while *n*_*y*_ is odd and, restricted to the interval 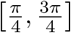, *n*_*x*_ has a central symmetry with respect to the points 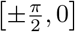, while *n*_*y*_ is symmetric with respect to the line 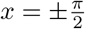, (see Figure 11a).

**Figure 11:**
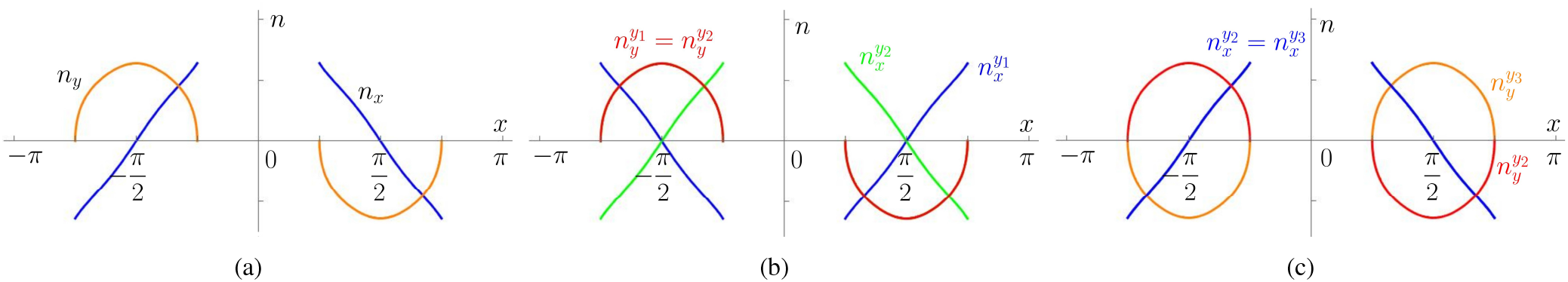
(a) Components of the normal vector for a point whose coordinates satisfy *y* = *y*_2_(*x*): in blue the function *n*_*x*_ and in orange the function *n*_*y*_. (b) Components of the normal vector for two points whose coordinates respectively satisfy *y* = *y*_1_(*x*) and *y* = *y*_2_(*x*) (in blue the function 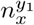, in red the function 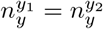 and in green the function 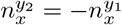). (c) Components of the normal vector for two points whose coordinates respectively satisfy *y* = *y*_2_(*x*) and *y* = *y*_3_(*x*) (in blue the function 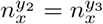, in red the function 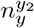 and in orange the function 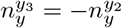).

Moreover, by symmetry, restricting our attention to the interval 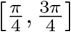, we have that *n*_*x*_ is monotonically decreasing while *n*_*y*_ is monotonically decreasing for 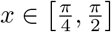, and monotonically increasing for 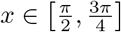.

To identify the points *A* and *B* that have opposite normal components, we exploit the just mentioned symmetries shown in Figure 11a. If we take for example 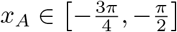 where *n*_*y*_ ≥ 0, then *x*_*B*_ must belong to 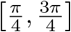 where *n*_*y*_ ≤ 0. Actually, it must be 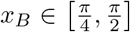 where *n*_*x*_ ≥ 0, because *n*_*x*_ ≤ 0 when 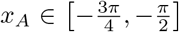 . Then, due to the just stated symmetries of *n*_*x*_ and *n*_*y*_

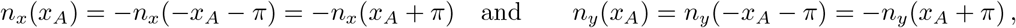

and so *x*_*B*_ = *x*_*A*_ + *π*.

Let us first examine the singular case characterized by *x*_*B*_ = *π/*2, corresponding to the red dot in Figure 9, and therefore *x*_*A*_ = −*π/*2. In this case 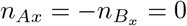 and

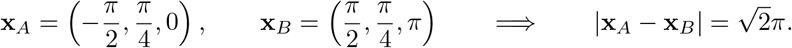

However, the two normal vectors **n**_*A*_ = (0, 1, 0) and **n**_*B*_ = (0, − 1, 0) do not belong to the same line, and the points do not belong to the same horizontal plane. So, *A* and *B* cannot be the endpoints of the diameter of the ball (see Figure 12a).

**Figure 12:**
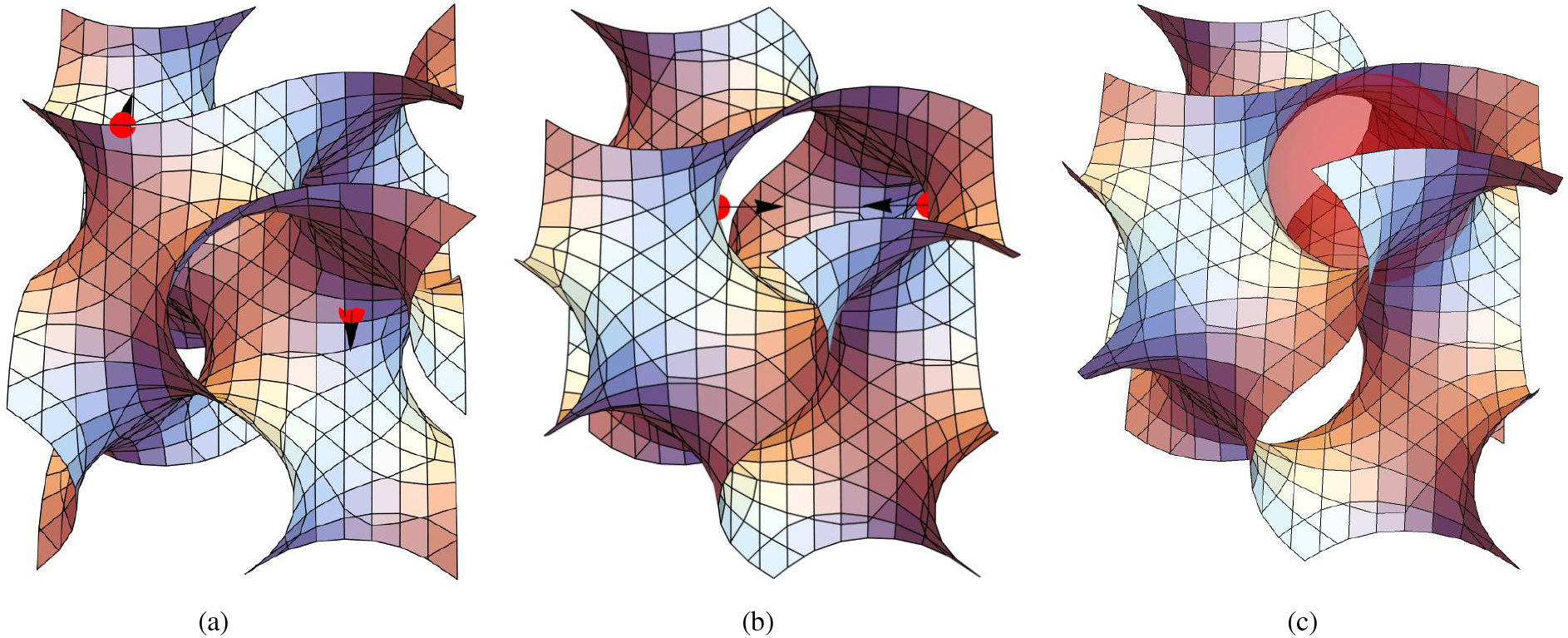
Couples of points with opposite normals with both points belonging to the same fold. In (a) 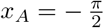 and 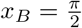 the normals are not aligned with the line joining *A* and *B*. In (b) 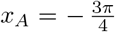 and 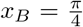 the normals are directly opposite, but as shown in (c) the pair is not admissible because the sphere interpenetrates the surface, as put in evidence by the shaded red area.

Noticing that sin(*x*_*B*_ − *π*) = − sin *x*_*B*_ and therefore *β*(*x*_*B*_ − *π*) = *β*(*x*_*B*_), for the non-singular cases with *x*_*B*_ *≠ π/*2 we can write

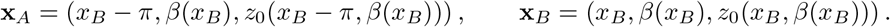

Thus

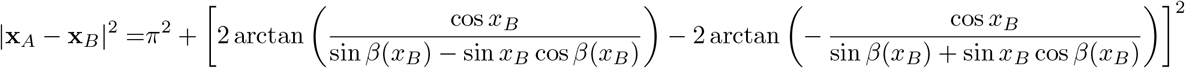

which is obviously minimized by |**x**_*A*_ − **x**_*B*_ | = *π* when the square parentheses vanish, i.e., when the arguments of the arctangents are equal, if this is possible. Actually, it can be readily checked that this occurs if and only if sin *β*(*x*_*B*_) = 0, that is, recalling (6.8), when 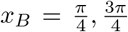. The other corresponding coordinates are then 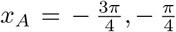 and *y*_*A*_ = *y*_*B*_ = 0, 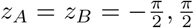, with **n**_*A*_ = (1, 0, 0) and **n**_*B*_ = (−1, 0, 0), see Figure 12b. However, checking the maximum curvature in both points one has 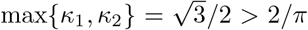. So, interpenetration holds and this solution is not admissible (see Figure 12c). This also hints at the fact that there are pairs of points with directly opposite normals and a distance smaller than *π*. In the following the diameter *D*_1_ = *π* is then used as a reference to exclude the cases leading to larger spheres.

In conclusion, this first case does not lead to the identification of admissible pairs of points.

#### 6.1.2 Projections on Different *y*-functions

A procedure similar to the one explained in the previous section can be applied when the projections of *A* and *B* on the plane *z* = 0 lie on different *y*-functions. In order to clarify the different situations a superscript *y*_*i*_ is added to the components of **n** to put in evidence which part of the boundary *y*_*i*_ the point belongs to. Let us consider the different cases:

1. One point projection lies on *y*_1_ and one on *y*_2_, e.g. *y*_*A*_ = *y*_1_(*x*_*A*_) = *π* − *β*(*x*_*A*_) and *y*_*B*_ = *y*_2_(*x*_*B*_) = *β*(*x*_*B*_): we obtain 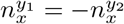 and 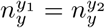, as shown in Figure 11b. If we take, for example, 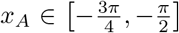 where 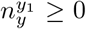, then *x*_*B*_ must belong to 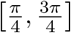 where 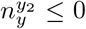. Actually, it must be 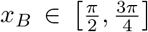 where 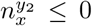, because 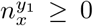 when 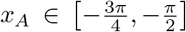. Then, due to the just stated symmetries of *n*_*x*_ and *n*_*y*_

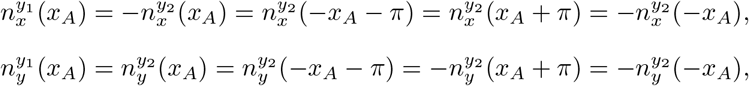

and so *x*_*B*_ = −*x*_*A*_ and

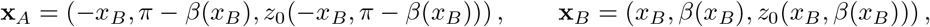

where we exploited the fact that *β*(−*x*_*B*_) = *β*(*x*_*B*_). Then, since *z*_0_(−*x*_*B*_, *π* − *β*(*x*_*B*_)) = *z*_0_(*x*_*B*_, *β*(*x*_*B*_)),

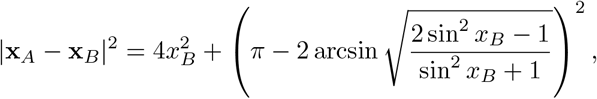

that is minimized by 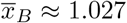. Then 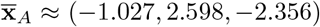 and 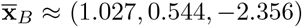, and 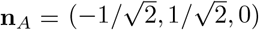, and 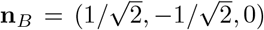, as shown in Figure 13a,b. Their distance is then *D*_2_ ≈ 2.904 *< D*_1_ = *π*, but checking for the maximal sectional curvature, we have that max {*κ*_1_, *κ*_2_ } ≈ 0.772 *>* 0.689 ≈ 2*/D*_2_. So, the tangent sphere would intersect the surface, as shown in Figure 13c, and therefore the pair of points is not admissible.
2. One point projection lies on *y*_2_ and one on *y*_3_, e.g. *y*_*A*_ = *y*_3_(*x*_*A*_) = −*β*(*x*_*A*_) and *y*_*B*_ = *y*_2_(*x*_*B*_) = *β*(*x*_*B*_): we obtain 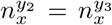 and 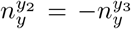, as one can see in Figure 11c. Then, proceeding as before one finds that *x*_*B*_ = *π* − *x*_*A*_. The distance between the two points is then minimized by 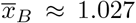. However, the points are 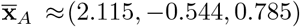 and 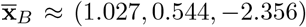 they have distance greater than *D*_1_ and they do not lie on the same horizontal plane, thus the couple is not admissible and is symmetrical to the case in Figure 12a.

**Figure 13:**
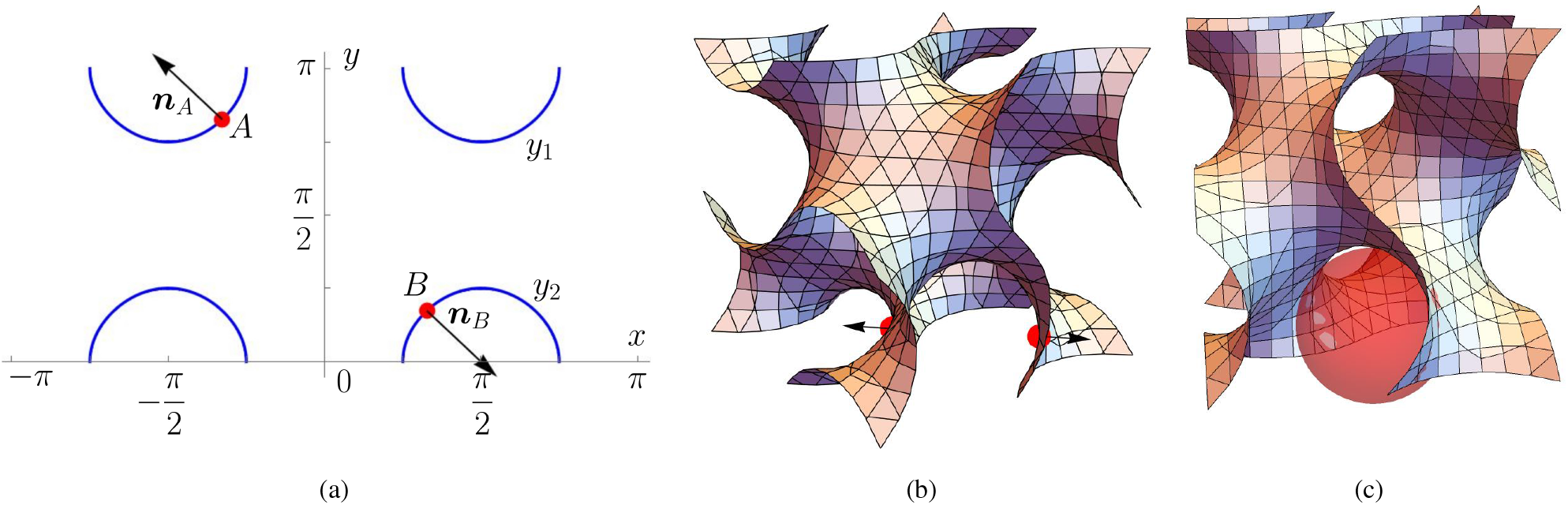
Couple of points with distance *D*_2_ and opposite normal vectors. Projections of *A* and *B* in a horizontal plane: (a) the projection of point *A* lies on *y*_1_(*x*) and that of *B* on *y*_2_(*x*) and their normal vectors have vanishing *z*-component. (b) The points *A* and *B* on the gyroid and (c) the corresponding sphere with interpenetration of matter with the surface.

Regarding the other cases, one can notice that, since *z*_*A*_ = *z*_*B*_, to evaluate the distance between the pairs of points, it is sufficient to consider the *x* and *y* coordinates. However, the distance in the plane between *y*_1_ and *y*_3_ or *y*_4_ is clearly greater than *D*_2_, and by symmetry, the case with *y*_3_ and *y*_4_ will give a couple of points with distance *D*_2_. All the other cases follow by symmetry. In conclusion, also this second case only leads to non-admissible pairs of points.

### 6.2 Points with opposite normal vectors: two points on different folds

First of all, we remark that, due to the invariance under cyclic permutations of the axes, the cases with *n*_*x*_ = 0 and *n*_*y*_ = 0 give results identical to those with *n*_*z*_ = 0 discussed in the previous subsection. In particular, there are no admissible pairs of contact points having a vanishing component of the normal. Thus, we can here reduce our study to the case where neither *n*_*x*_ nor *n*_*y*_ nor *n*_*z*_ vanish for both points.

Therefore, we have to look for two points *A* and *B* belonging respectively to the lower and the upper fold, that is *z*_*A*_ = *z*_−_(*x*_*A*_, *y*_*A*_) and *z*_*B*_ = *z*_+_(*x*_*B*_, *y*_*B*_).

Figure 14 reports the domains where 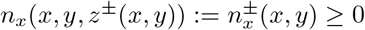 and 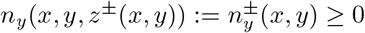 for the two folds, as determined in Appendix B.

**Figure 14:**
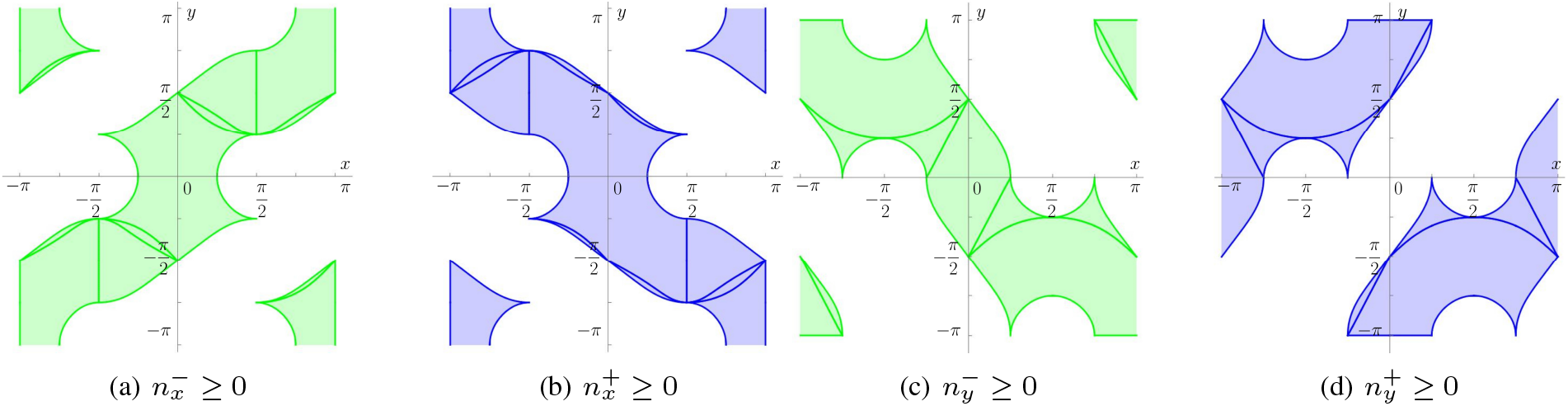
The domains of positivity of (a) 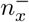 (b) 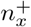 (c) 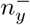 and (d) 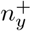..

In order to identify the points with directly opposite normals and in particular such that 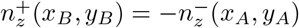, it is useful to give a closer look at the functions 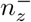 and 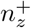 drawn in Figure 10 and at their symmetries. First of all, 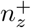 is always negative, while 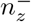 is always positive, and 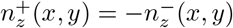.

Then, focusing on 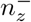, it results that

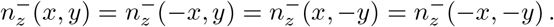

Moreover, taking for instance the set 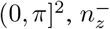 is symmetrical with respect to the lines 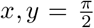 and to the points 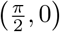.

The same holds for the other quadrants.

The function 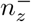 presents global maxima when *x* = 0, ±*π* and 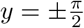, such that

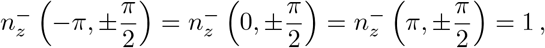

and saddle points when *x, y* = 0, *±π* and 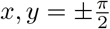 with all combinations of signs and

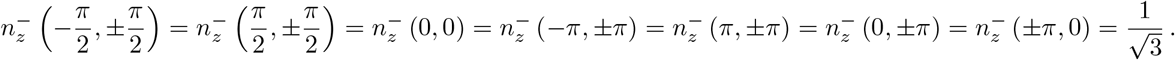

Thus, if 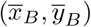 is a candidate point on the blue fold in Figure 10, (*x*_*A*_, *y*_*A*_) must belong to a level set of 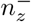 so that 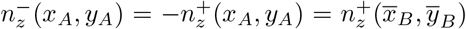 on the orange fold where the level sets are again curves that satisfy all the just mentioned symmetries (see again Figure 10).

Focusing on the other components of the normal, taking into account all the possible signs of the normal components, we have four possible cases:

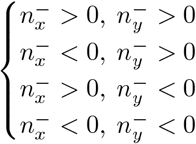

in (*x*_*A*_, *y*_*A*_) with the corresponding opposite signs for 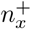 and 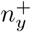 in (*x*_*B*_, *y*_*B*_). The couples of *x* and *y* coordinates of points satisfying these inequalities identify four domains that present a high symmetry with respect to each other. For this reason, we restrict our discussion to the first case, being those of the others completely similar. So, we refer to the plot in Figure 15, giving the domains where 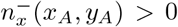 and 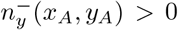 (in red) and 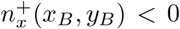 and 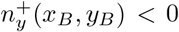 (in yellow). We remark that the case (*x*_*A*_, *y*_*A*_) = (*x*_*B*_, *y*_*B*_) was already excluded in the previous section because *n*_*z*_ would vanish.

**Figure 15:**
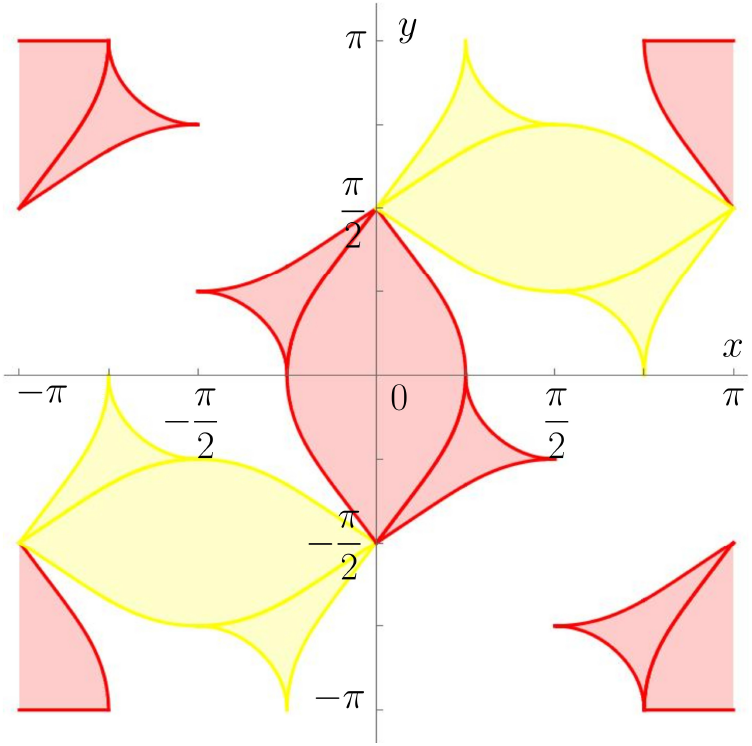
Domains where (*x*_*A*_, *y*_*A*_) is such that 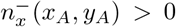 and 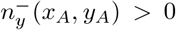 (red) and (*x*_*B*_, *y*_*B*_) is such that 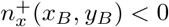 and 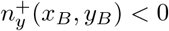 (yellow).

The study of the level sets of the normal components, that depends on four variables (*x*_*A*_, *y*_*A*_, *x*_*B*_, *y*_*B*_) at the moment independent, is not straightforward and requires a numerical support (discussed at the end of this section). In fact, as it can be realized looking at Figure 16, it is impossible to find an explicit analytic form for the level sets to identify all the possible couples of points with opposite normals. However, due to the high level of symmetry of triply periodic surfaces, we expect that the coordinates of the two points we are looking for will satisfy the symmetries characterizing the components of the normals 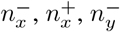 and 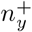 (see, respectively, Figure 16a (orange), 16a (blue), 16b (orange), 16b (blue)). Therefore, we start our analysis looking for them. Similarly to the symmetries of 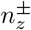, we have 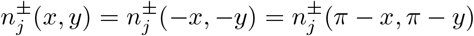 for both *j* = *x, y*. So, in Figure 15 we could restrict our attention to the central red domain and one of the yellow ones.

**Figure 16:**
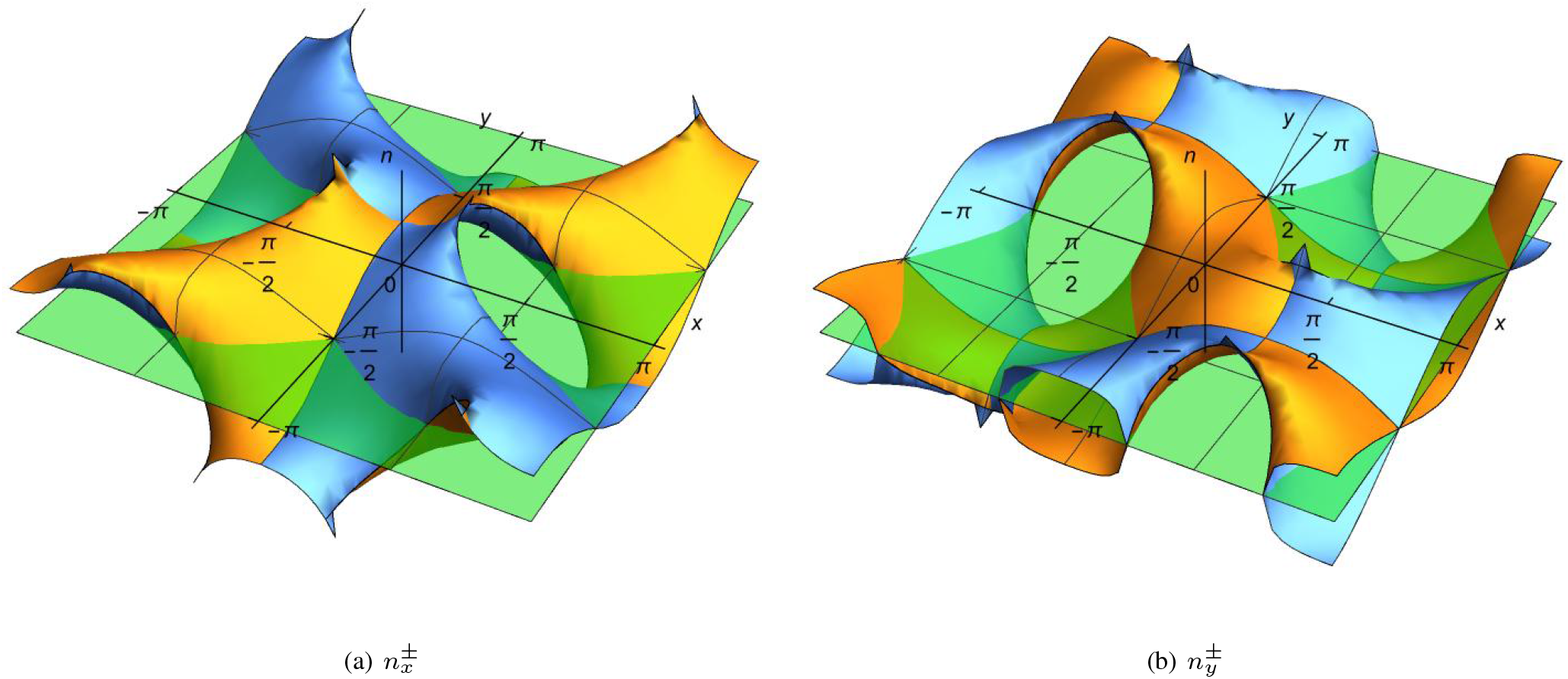
(a) The functions 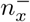 and 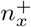. (b) The functions 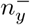 and 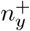. In orange, the quantities corresponding to the fold *z*_−_, in blue the ones corresponding to *z*_+_. In green, the plane *n* = 0.

Moreover, 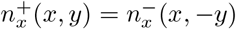 and 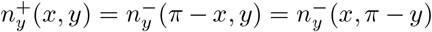, and the same, of course, holds for their level sets. Thus, the two points must belong to the same level set of 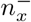 and 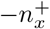 and to the same level set of 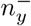 and 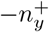. We remark that:

- the level sets of 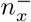 and the level sets of 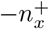 are symmetric with respect to the lines 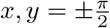
- the level sets of 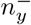 and the level sets of 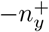 are symmetric with respect to the *x* and *y* axis.

Regarding the symmetries of the level sets of the *x*-component, the two points (*x*_*A*_, *y*_*A*_) and (*x*_*B*_, *y*_*B*_) can be symmetric with respect to the lines 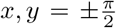 or can satisfy (*x*_*A*_, *y*_*A*_) = (*x*_*B*_ − *π*, − *y*_*B*_), or (*x*_*A*_, *y*_*A*_) = (− *x*_*B*_, *y*_*B*_ − *π*), see red dots in Figure 17. Regarding the symmetries of the level sets of the *y*-component, the two points *A* and *B* can be symmetric with respect to the lines *x, y* = 0 or can satisfy (*x*_*A*_, *y*_*A*_) = (*x*_*B*_ − *π, π* − *y*_*B*_), or (*x*_*A*_, *y*_*A*_) = (*π* − *x*_*B*_, *y*_*B*_ − *π*), see blue dots in Figure 17. Summing up, given a point (*x*_*B*_, *y*_*B*_) we have four possible symmetries as regards *n*_*x*_, four for *n*_*y*_ and sixteen for *n*_*z*_.

**Figure 17:**
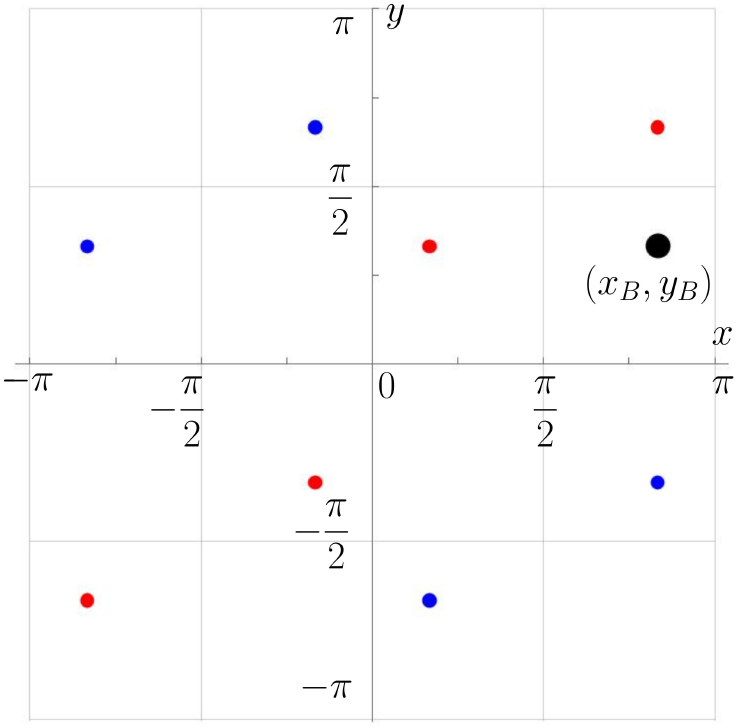
Starting from the reference point (*x*_*B*_, *y*_*B*_), labeled in black, in red, the possible positions of (*x*_*A*_, *y*_*A*_) such that *n*_*x*_(*x*_*A*_, *y*_*A*_) = −*n*_*x*_(*x*_*B*_, *y*_*B*_) when we consider the symmetries of the function *n*_*x*_. In blue, the same for (*x*_*A*_, *y*_*A*_) such that *n*_*y*_(*x*_*A*_, *y*_*A*_) = −*n*_*y*_(*x*_*B*_, *y*_*B*_). One can notice that for any choice of *x*_*B*_, *y*_*B*_ *>* 0, there are no points such that simultaneously *n*_*x*_(*x*_*A*_, *y*_*A*_) = −*n*_*x*_(*x*_*B*_, *y*_*B*_) and *n*_*y*_(*x*_*A*_, *y*_*A*_) = −*n*_*y*_(*x*_*B*_, *y*_*B*_).

However, as it can be seen in Figure 17, once given the coordinates (*x*_*B*_, *y*_*B*_), in general the intersection of the possible coordinates for (*x*_*A*_, *y*_*A*_) respecting the symmetries of *n*_*x*_ and *n*_*y*_ provides an empty set of couple of points with all the three components of the normal vectors equal and opposite in our domains.

Regarding instead the critical points of 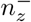 lying in the red and yellow domains, computing the quantities for the lower and upper folds depending on the red or yellow domain, we have either

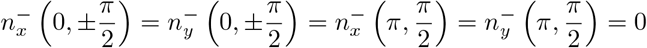

or

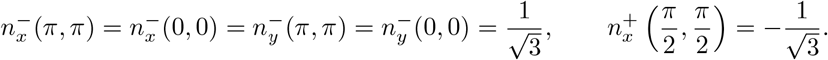

The former have *n*_*y*_ = 0 and have been already discussed in the previous section with the conclusion that they lead to non-admissible pairs (see the comment at the end of the section).

Regarding the latter, by symmetry we can take as an example the points **x**_*A*_ = (0, 0, 0) and 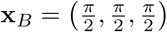, with normal vectors 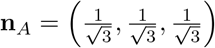 and 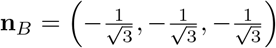. The two points, shown in Figure 18a, thus identify the diameter of the largest sphere moving through the gyroid without getting stuck; they have distance 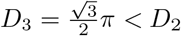.

**Figure 18:**
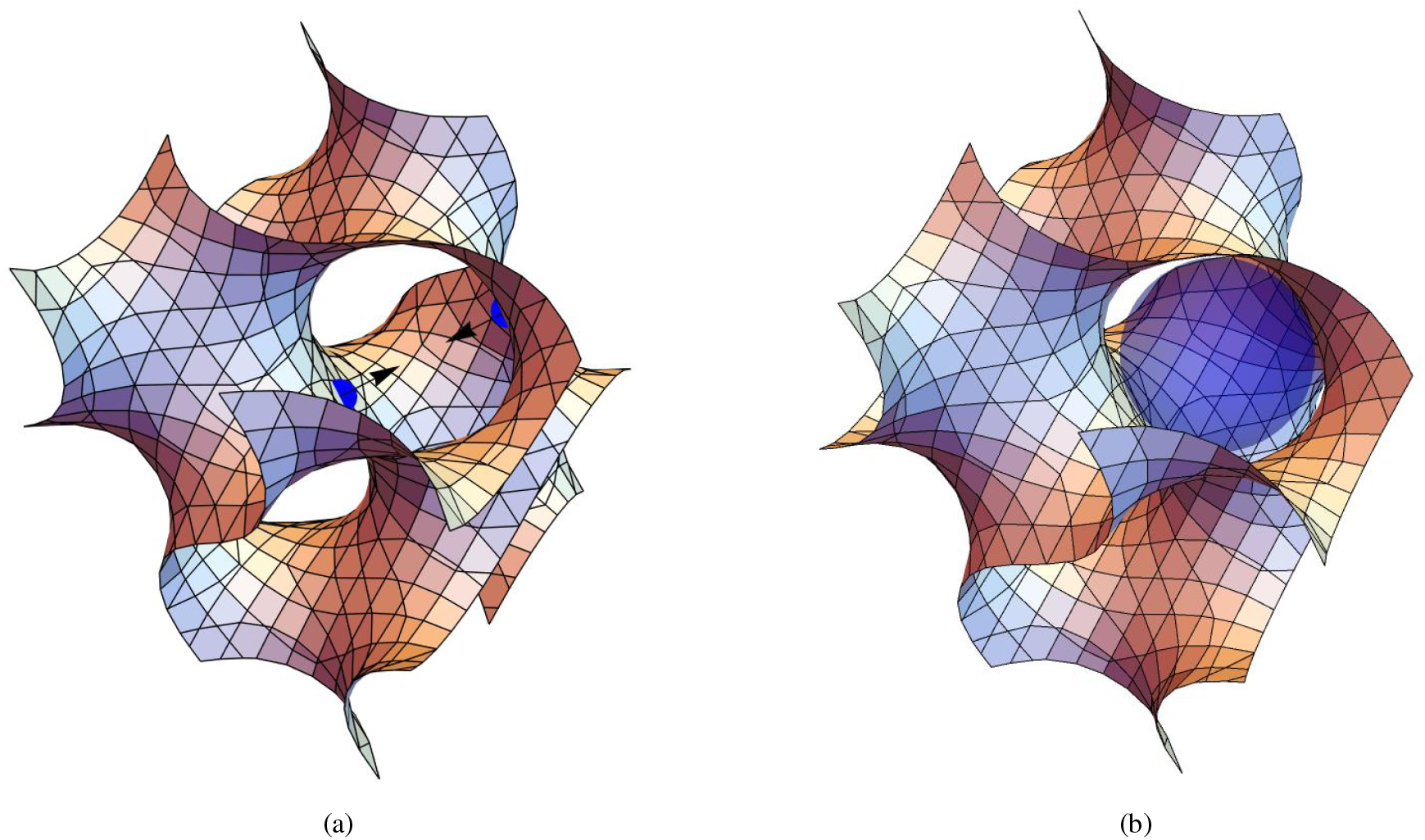
(a) The couple of points **x**_*A*_ = (0, 0, 0), 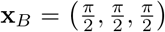 that give the diameter *D*_3_ of the biggest sphere moving in the structure. (b) The sphere with diameter *D*_3_ passing in the gyroid that does not present interpenetration of matter.

Computing the principal curvatures of the surface at these two points from the gaussian and the mean curvature, it can be verified that locally the sphere does not interpenetrate the surface. Actually, there *κ*_1_ = *κ*_2_ = 0 and thus the surface is locally flat, so the sphere surely does not intersect the surface around these points, as one can see in Figure 18b, since the maximal curvature of the surface section is lower than the curvature of the tangent sphere which will pass through the gyroid.

As regards all the other points belonging to the same level sets but not satisfying the symmetries treated above, one can verify through a simple numerical simulation that there are no other points satisfying the requirement of having opposite normals, and the admissible couples with minimal distance are precisely all the couples symmetrical to our solution.

In conclusion, using the diameter of the sphere as reference, the dimension of the periodic cell of the gyroid must be

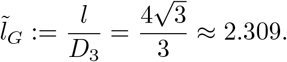

## 7 Conclusions

With the aim of identifying what are the geometrical characteristics that a biomimetic scaffold must have in order to facilitate cell migration and to foster then its rapid re-population, we studied several triply-periodic structures. The starting idea is that during cell migration the cell nucleus, taken as a sphere, must not deform during motion and that the cell membrane need to adhere to the walls of the scaffold to be able to exert traction forces.

Thanks to the symmetry and periodicity of the structures studied, we find, for instance, that the ratio between the size of the periodic cell of the structure and the diameter of the nucleus of the cell that need to comfortably crawl in the scaffold is 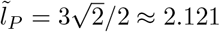 in the case of a Schwarz P-surface, 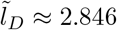 in the case of a Schwarz D-surface and 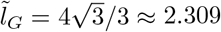 in the case of a gyroid.

Always using as a reference length the size of the nucleus, the ratio of the surface area of the structure (that the cell can use to adhere) with respect to the area of the cell nucleus is 𝒜_*P*_ ≈ 3.38 for the Schwarz P-surface, 𝒜_*D*_ ≈ 3.74 for the Schwarz D-surface and 𝒜_*G*_ ≈ 10.3 for the gyroid, suggesting that more complicated and twisted surfaces as the gyroid can provide a much larger surface to which cells can adhere through the cytoskeleton. Indeed, the additive manufacturing of 3D bio-mimetic scaffolds exploits intricate shapes.

The strategy applied in this work can naturally be extended to other surfaces, both analytically, if the surface presents enough degrees of symmetry, or numerically. A forthcoming paper will instead focus on the analysis of the possible paths that the cell can follow into these structures, in view of a first classification of these five lattices in terms of their optimality, i.e. the ratio between the distance between the endpoints of the path and its length. Another research direction of big interest could be to numerically test 3D models of cell migration such as those using CPM or phase field models such as [18] in such geometries.

## Acknowledgments

The authors thank Alessandro Giammarini, Luca Lussardi, Alfredo Marzocchi and Fabio Vicini for fruitful discussions. CL is supported by the PNRR M1C2 - CDS000965 Silicon Valley 8.0 (CUP C17J23000030001). CL and LP were also supported by the MICS (Made in Italy - Circular and Sustainable) Extended Partnership and received funding from the European Union Next-Generation EU (PIANO NAZIONALE DI RIPRESA E RESILIENZA (PNRR) - MISSIONE 4 COMPONENTE 2, INVESTIMENTO 1.3 - D.D. 1551.11-10-2022, PE00000004). This work was partially conducted also under the National Plan for Complementary Investments to the NRRP, project “D34H-Digital Driven Diagnostics, prognostics and therapeutics for sustainable Health care” (project code: PNC0000001), Spoke 4 funded by the Italian Ministry of University and Research. CL and LP also acknowledges the support of Gruppo Nazionale per la Fisica Matematica (GNFM) of Istituto Nazionale di Alta Matematica (INdAM); CL was supported through the “INdAM - GNFM Project”, codice CUP #E5324001950001#.

## A Complete computations with the analytical procedure for two points with op-posite normals on the Schwarz P-surface

We proceed here with the technique explained in Subsection 3.1 that allows to find the smallest passage inside the cell. *Case I: the two points belong to the same fold*.

We choose to work on the fold with *z* ≥ 0, so both points *A* and *B* with opposite normals we are looking for will have 0 ≤ *z* ≤ *π*. Being the third component of the normal vector always negative for such points and requiring opposite components for them, *n*_*z*_ must vanish, i.e., either *z*_*A*_ = *z*_*B*_ = 0 or *z*_*A*_ = *z*_*B*_ = *π*. Because of symmetry, without loss of generality we will also assume that *x*_*A*_, *y*_*A*_ ≥ 0, which also implies that *x*_*B*_, *y*_*B*_ ≤ 0 because of the opposite signs of the normals.

- If *z* = *π* for both points, the equality *n*_*z*_ = 0 implies that the points lie on *y*_1_(*x*) or *y*_2_(*x*) on the central yellow curves in Figure 6. Assuming that they both lie, for example, on *y*_2_, we have

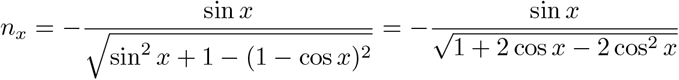

that is an odd function, positive for *x <* 0, negative for *x >* 0, and strictly decreasing since

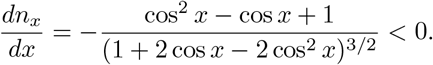 This implies that to have opposite *x*-components of the normal for two values of *x*, one must have *x*_*B*_ = −*x*_*A*_. Operating in a similar way on *n*_*y*_ would give *y*_*B*_ = −*y*_*A*_. So,

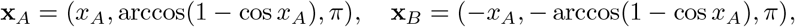

with *x*_*A*_ ∈ [0, *π/*2]. Then, we want to minimize

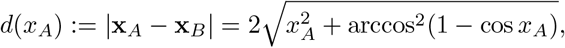

in [0, *π/*2]. Inside this interval, *d*(*x*_*A*_) is minimized when

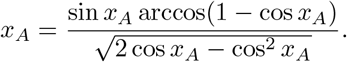 A graphical study shows that the above equation has only one root given by 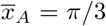, yielding 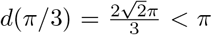 identifying the minimum of *d*. Then, the diameter of the ball is 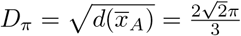, that confirms the value obtained in Subsection 4.1, with **x**_*A*_ = (*π/*3, *π/*3, *π*) and **x**_*B*_ = (−*π/*3, −*π/*3, *π*) and normal vectors 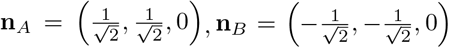, (see Figures 3a,c and blue points in Figure 5a). On the other hand, it can be readily observed that *d*(0) = *d*(*π/*2) = *π*, that completes our treatment of this case, since when a point belongs to *y*_1_ and one to *y*_2_, with an analogous reasoning, that gives |**x**_*A*_ − **x**_**B**_ |= *d*(*π/*2) = *π > D*_*π*_ (see Figures 3b and red points in Figure 5a) and thus do not determine the smallest constriction.
- If *z* = 0 for both points, the equality *n*_*z*_ = 0 implies that the points lie on *y*_3_(*x*) or *y*_4_(*x*), i.e. the blue curves in Figure 6. But with some sign changes, one can repeat the same steps as above to find again that *x*_*B*_ = − *x*_*A*_ and *y*_*B*_ = − *y*_*A*_, so that

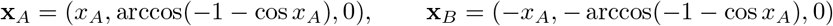

and

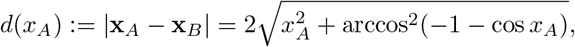

defined in [*π/*2, *π*]. The minimum is then obtained when *x*_*A*_ = 2*π/*3 with the points of contact given by (see Figures 4a,b,c (in red) and Figure 5b)

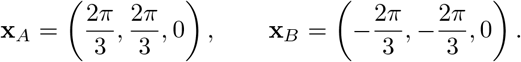 The minimum distance when *z* = 0 is then 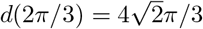 which is larger than the one found when *z* = *π*;
- both points lie along *y*_4_ (or *y*_3_): then, recalling *z* = 0, the normal vectors for both points read

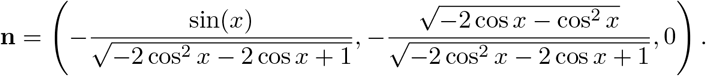 The *y*-component is negative, so to have two opposite components we need it to van ish: thu s, as before, *x*_*A,B*_ = *± π/*2 and in particular *x*_*A*_ = − *π/*2 and *x*_*B*_ = *π/*2; then *n*_*Ax*_ = − *n*_*Bx*_ and finally 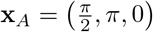 and 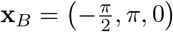 that have distance *π > D*_*π*_: it is a symmetric case of ones of the above.

*Case II: the two points belong to different folds*.

The only interesting case has been treated in Section 4.2.2. The other cases imply that at least *n*_*x*_ or *n*_*y*_ vanishes, but by symmetry these cases are equivalent to the one already treated with *n*_*z*_ = 0 and they both lead to couples of points with distance greater than *π* or lying in distinct planes, in contradiction with Remark 3.2.

## B Study of the signs of 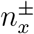 and 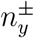 for the gyroid

The aim of this appendix is to identify where 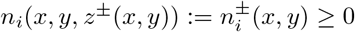 for *i* = *x, y* for the two folds.

### B.1 Sign of 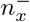.

With the same simplifications used at the beginning of Subsection 6.1, we obtain that

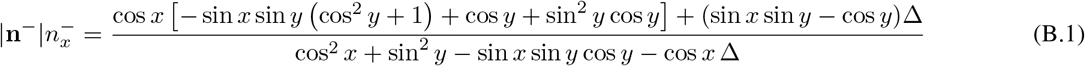

and the denominator is a positive quantity for the points on the two folds (see Appendix C). So, it is enough to study the inequality

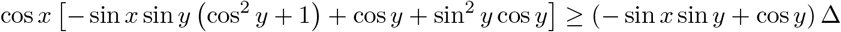

in the domain of the gyroid, which assures that the square root Δ is well-defined. To study this irrational inequality, we can analyze three different cases.

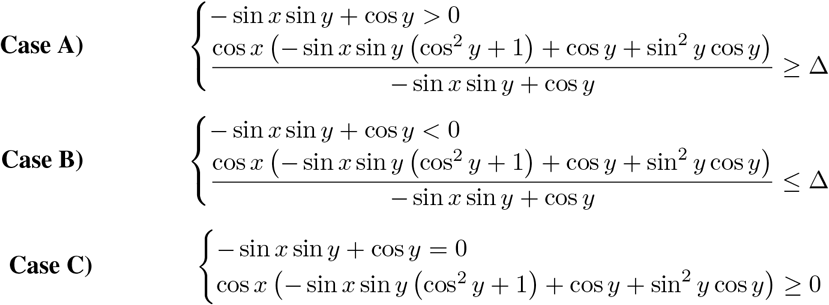

**Case A)**

The first inequality can be solved in terms of *y* and holds if

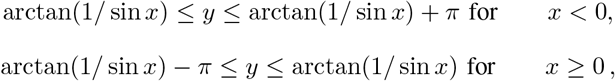

that is, the red domain in Figure 19a. The second inequality requires to study

**Figure 19:**
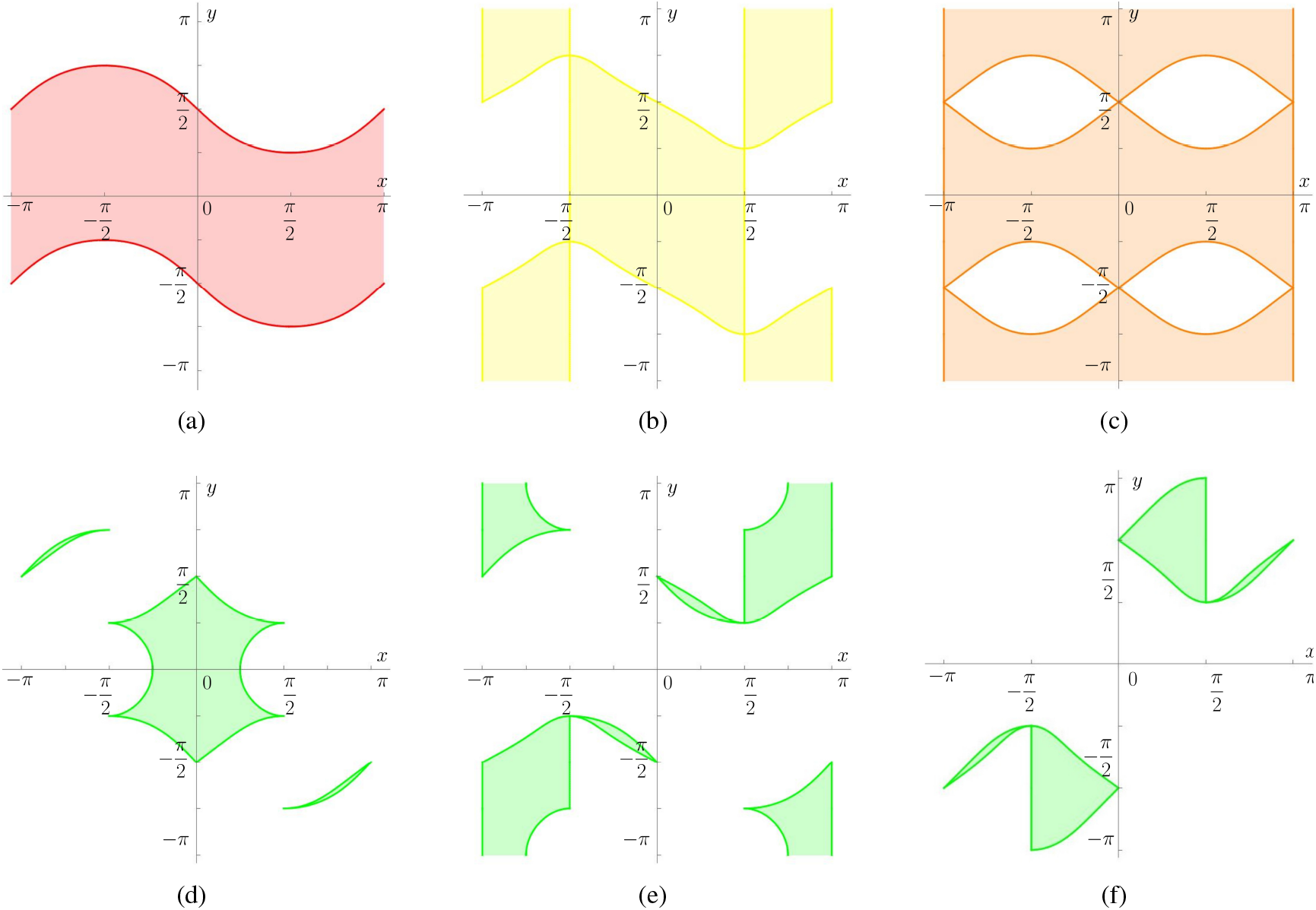
In the top row, the three distinct domains obtained from Case A) in the study of 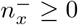. (d) The set obtained from Case A), as intersection of the red, yellow and orange domains in (a), (b), (c). In (e) and (f), respectively, the sets obtained from the first inequality of Case B) and the system in Case B) for the study of 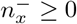.

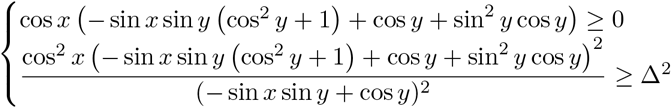

where the first inequality can be decomposed into two terms and, in particular, the second can be analytically studied in terms of sin *x*. This leads to the yellow domain in Figure 19b. The second one instead with standard simplifications and trigonometric addition formulas can be reduced to

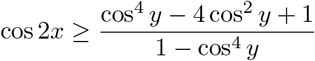

that can be inverted and studied analytically, to obtain the orange domain in Figure 19c.

The intersection of the just presented three domains and the domain of the gyroid gives for case A) the green set in Figure 19d.

**Case B)**

The first inequality clearly gives the complementary domain to the one in 19a. The second inequality requires to study

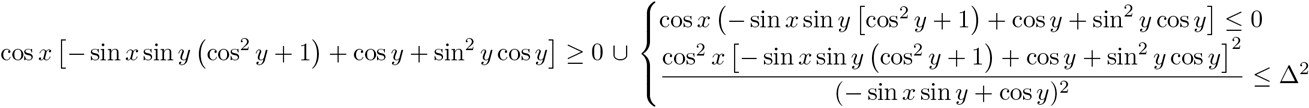

where the analysis can be easily linked back to the previously obtained domains. The two sub-cases, respectively, lead to the domains in Figure 19e,f.

**Case C)**

This case simply provides that the boundary of the red domain in Figure 19a that lies in the yellow domain in Figure 19b is taken into account in the green domain in Figure 14a, as drawn.

The union of the three cases gives the domain where 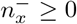, which is shown in Figure 14a.

### B.2 Sign of 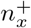.

Similarly, we can restrict our attention to the numerator of 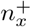, and study the inequality

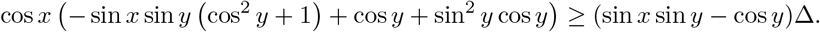

As before, we can distinguish different cases and obtain computations and domains analogous or complementary to those obtained before. The domain of positivity of 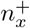 is shown in Figure 14b.

### B.3 Sign of 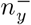.

Similarly to before, we have to study the inequality

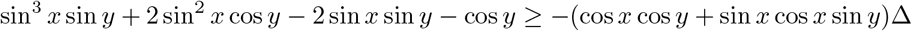

in the domain of the gyroid. We can distinguish three cases.

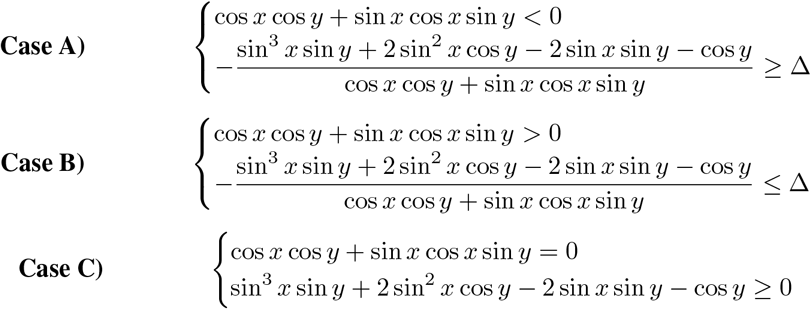

**Case A)**

In the domain of the gyroid, we study the system

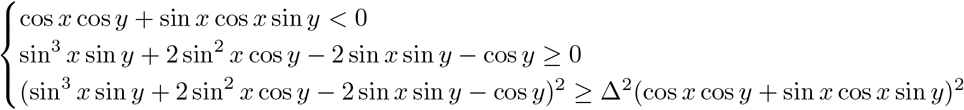

As before, all these inequalities can be analytically solved: the first two can be solved in terms of tan *y* and the third as in the previous cases by simplifications and trigonometric addition formulas, obtaining an inequality for cos *y*. The three inequalities, respectively, give the three domains in Figure 20a,b,c. In the domain of the gyroid, the system is satisfied by the green domain in Figure 20d.

**Figure 20:**
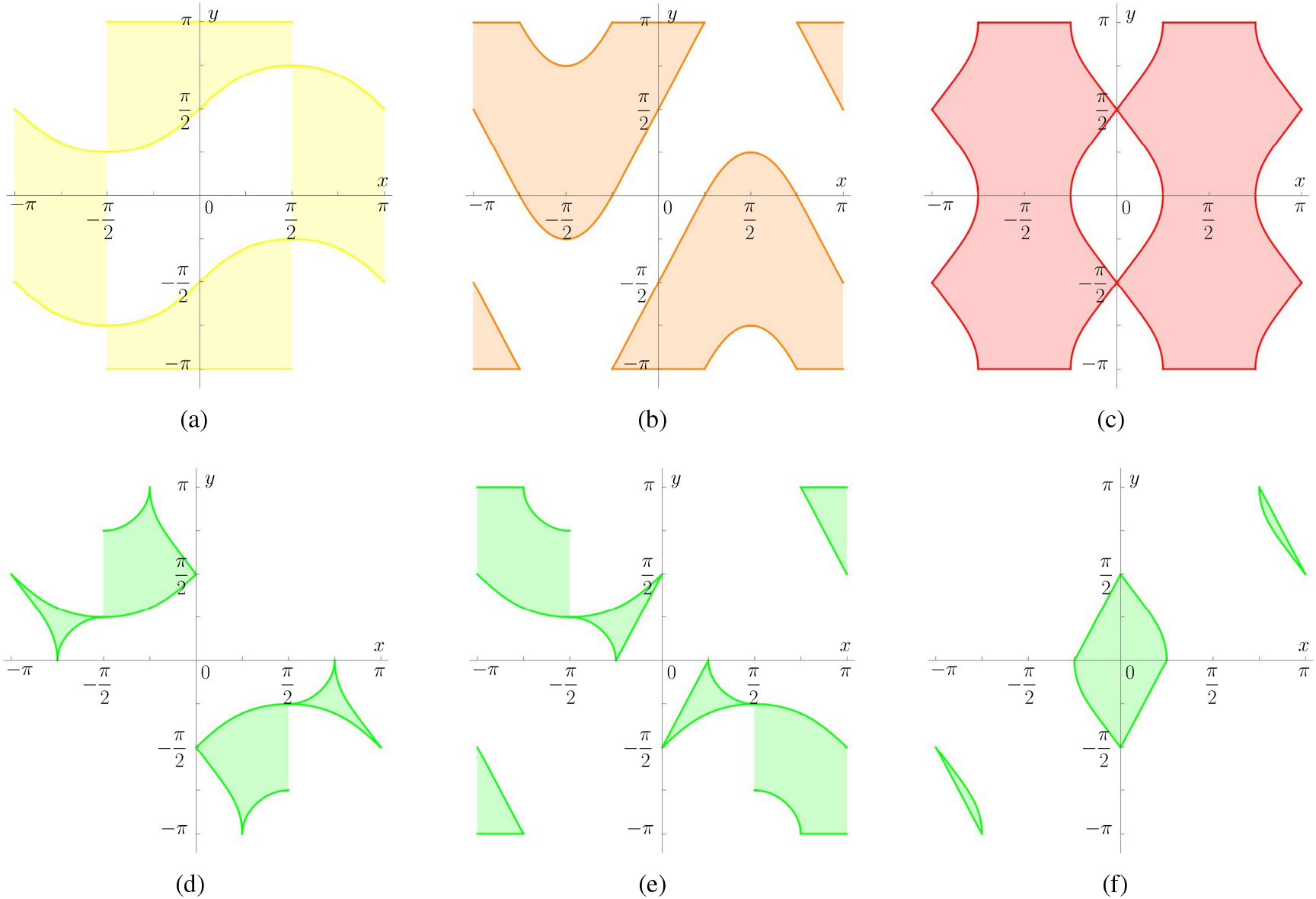
In the top row, the three distinct domains obtained from Case A) in the study of 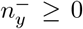. (d) The set obtained from Case A), as the intersection of the yellow, orange and yellow domains in (a), (b), (c). In (e) and (f), respectively, the sets obtained from the first inequality of Case B) and the system in Case B) for the study of 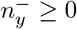.

**Case B)**

In the domain of the gyroid, we analyze the system

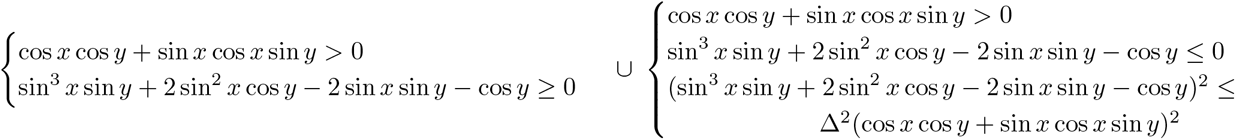

where the analysis can be easily reconduct to the domains previously obtained. The two sub-cases respectively lead to domains in Figure 20e,f.

**Case C)**

This case simply provides that the boundary of the yellow domain in Figure 20a that lies in the orange domain in Figure 20b is taken into account in the green domain in Figure 14c, as drawn.

The union of the three cases gives the domain where 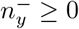, that is drawn in Figure 14c.

### B.4 Sign of 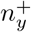

Similarly to the cases above, we can restrict our attention to the numerator of 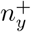 and to study the inequality

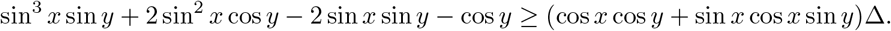

As before, we can distinguish different cases and obtain computations and domains analogous or complementary to those obtained before. The domain of positivity of 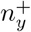 is drawn in Figure 14d.

## C Proof of the positivity of the denominator in (B.1)

We briefly show that

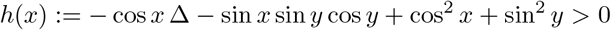

for both folds of the gyroid to complete the treatment of Appendix B. We thus analyze

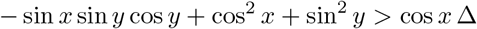

and we have three cases. We also notice that

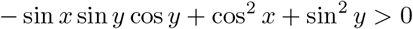

in all the domain of the gyroid apart from the points 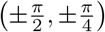 and 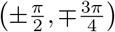 where the folds are not defined. One can carry out explicit computations (that we do not report here), since the expression can be rewritten as

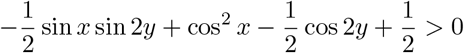

and solved in terms of *y*. We report here graphically in Figure 21 the positivity domain of this expression in red, compared with that of the gyroid in blue. The black lines represent the set of points where the two folds are not defined inside the blue domain.

**Figure 21:**
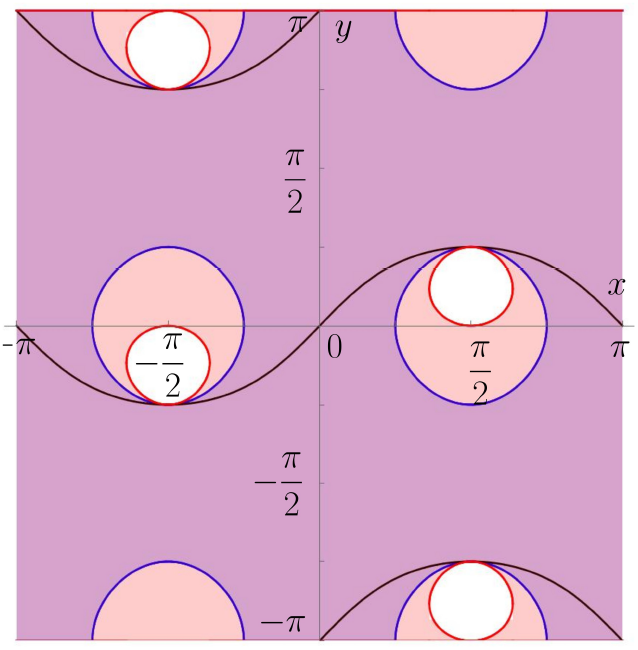
The domain of positivity of − sin *x* sin *y* cos *y* + cos^2^ *x* + sin^2^ *y* is in pink, i.e. outside the little circles, while the domain of the gyroid is the one outside the bigger circle. The intersection of the two domains give the purple domain, that assures that the positivity of the function in the domain of the gyroid.

**Case 1)** In the domain of the gyroid, we solve

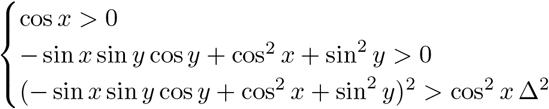

This can be easily solved since the second inequality is always verified in the domain of the gyroid where cos *x >* 0 and since the third equation can be simplified into

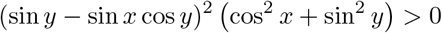

that is always positive and vanishes when sin *y* = sin *x* cos *y*, that is outside the domain of the folds (case *b* = *c*) or when cos^2^ *x* + sin^2^ *y* = 0, i.e. the points with *y* = 0, ±*π* and *x* = ±*π/*2, that are all outside the domain of the gyroid.

**Case 2)** In the domain of the gyroid, we solve

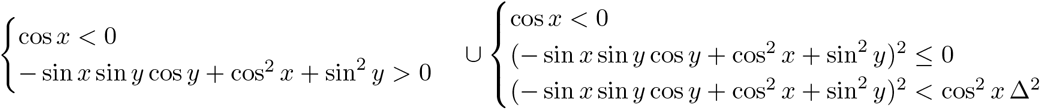

This can be easily solved since the first system is always true again because the second inequality is always verified in the domain of the gyroid where cos *x <* 0.

**Case 3)** In the domain of the gyroid, we solve

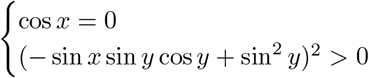

that again is true by the above considerations since − sin *x* sin *y* cos *y* + sin^2^ *y* vanishes only for points 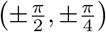 and 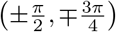 on the black curves in Figure 21 on which the folds are not defined.

## References

[1] V. te Boekhorst, L. Preziosi, and P. Friedl. Plasticity of cell migration in vivo and in silico. Annual Review of Cell and Developmental Biology 32 (2016), pp. 491–526.

[2] S. J. P. Callens, R. J. C. Uyttendaele, L. E. Fratila-Apachitei, and A. A. Zadpoor. Substrate curvature as a cue to guide spatiotemporal cell and tissue organization. Biomaterials 232 (2020), p. 119739.

[3] A. P. G. Castro, T. Pires, J. E. Santos, B. P. Gouveia, and P. R. Fernandes. Permeability versus Design in TPMS Scaffolds. Materials 12:8 (2019). DOI: 10.3390/ma12081313.

[4] J. Chen, D. Weihs, M. van Dijk, and F. J. Vermolen. A phenomenological model for cell and nucleus deformation during cancer metastasis. Biomechanics and Modeling in Mechanobiology 17 (2018), pp. 1429–1450.

[5] M. P. Do Carmo. Differential geometry of curves and surfaces: revised and updated second edition. Courier Dover Publications, 2016.

[6] A. D. Doyle, F. W. Wang, K. Matsumoto, and K. M. Yamada. One-dimensional topography underlies three-dimensional fibrillar cell migration. Journal of Cell Biology 184 (2009), pp. 481–490.

[7] P. Friedl and E. B. Bröcker. The biology of cell locomotion within three-dimensional extracellular matrix. Cellular and Molecular Life Sciences 57 (2000), pp. 41–64.

[8] P. Friedl and K. Wolf. Tumour-cell invasion and migration: diversity and escape mechanisms. Nature Reviews Cancer 3 (2003), pp. 362–374.

[9] C. Giverso, A. Arduino, and L. Preziosi. How nucleus mechanics and ECM microstructure influence the invasion of single cells and multicellular aggregates. Bulletin of Mathematical Biology 80 (2018), pp. 1017–1045. DOI: 10.1007/s11538-017-0262-9.

[10] C. Giverso, A. Grillo, and L. Preziosi. Influence of nuclear deformability on cell entry into cylindrical structures. Biomechanics and Modeling in Mechanobiology 13 (2014), pp. 481–502. DOI: 10.1007/s10237-013-0510-3.

[11] C. Giverso, G. Jankowiak, L. Preziosi, and C. Schmeiser. The influence of nucleus mechanics in modelling adhesion-independent cell migration in structured and confined environments. Bulletin of Mathematical Biology 85 (2023), p. 88. DOI: 10.1007/s11538-023-01187-8.

[12] B. A. Harley, H. Kim, M. H. Zaman, I. V. Yannas, D. A. Lauffenburger, and L. J. Gibson. Microarchitecture of three-dimensional scaffolds influences cell migration behavior via junction interactions. Biophysical Journal 95 (2008), pp. 4013–4024.

[13] R. M. Kuntz and W. M. Saltzman. Neutrophil motility in extracellular matrix gels: mesh size and adhesion affect speed of migration. Biophysical Journal 72 (1997), pp. 1472–1480.

[14] D. A. Lauffenburger and A. F. Horwitz. Cell migration: a physically integrated molecular process. Cell 84 (1996), pp. 359–369.

[15] L. Li, J. Shi, K. Zhang, L. Yang, F. Yu, L. Zhu, H. Liang, X. Wang, and Q. Jiang. Early osteointegration evaluation of porous Ti6Al4V scaffolds designed based on triply periodic minimal surface models. Journal of Orthopaedic Translation 19 (2019), pp. 94–105. DOI: 10.1016/j.jot.2019.03.003. URL: https://www.sciencedirect.com/science/article/pii/S2214031X18302493.

[16] C. Lonati and A. Marzocchi. On self-contact and non-interpenetration of elastic rods. Mathematics and Mechanics of Solids 30:2 (2025), pp. 450–469.

[17] O. Maxian, A. Mogilner, and W. Strychalski. Computational estimates of mechanical constraints on cell migration through the extracellular matrix. PLoS Comput Biol 16:8 (2020), e1008160.

[18] A. Moure and H. Gomez. Phase-Field Modeling of Individual and Collective Cell Migration: A. Moure, H. Gomez. Archives of Computational Methods in Engineering 28:2 (2021), pp. 311–344.

[19] S. R. Peyton and A. J. Putnam. Extracellular matrix rigidity governs smooth muscle cell motility in a biphasic fashion. Journal of Cellular Physiology 204 (2005), pp. 198–209.

[20] R. Pugliese and S. Graziosi. Biomimetic scaffolds using triply periodic minimal surface-based porous structures for biomedical applications. SLAS Technology 28:3 (2023), pp. 165–182. DOI: 10.1016/j.slast.2023.04.004.

[21] C. G. Rolli, T. Seufferlein, R. Kemkemer, and J. P. Spatz. Impact of tumor cell cytoskeleton organization on invasiveness and migration: a microchannel-based approach. PLoS ONE 5 (2010), e8726.

[22] F. Sabeh, R. Shimizu-Hirota, and S. J. Weiss. Protease-dependent versus-independent cancer cell invasion programs: three-dimensional amoeboid movement revisited. Journal of Cell Biology 185 (2009), pp. 11–19.

[23] M. Scianna and L. Preziosi. A Cellular Potts Model analyzing cancer cell migration across constraining pillar arrays. Axioms 10 (2021), p. 32.

[24] M. Scianna and L. Preziosi. Modeling the influence of nucleus elasticity on cell invasion in fiber networks and microchannels. Journal of Theoretical Biology 317 (2013), pp. 394–406. DOI: 10.1016/j.jtbi.2012.11.003.

[25] M. Scianna, L. Preziosi, and K. Wolf. A Cellular Potts Model simulating cell migration on and in matrix environments. Mathematical Biosciences and Engineering 10 (2013), pp. 235–261. DOI: 10.3934/mbe.2013.10.235.

[26] X. Trepat, Z. Chen, and K. Jacobson. Cell migration. Comprehensive Physiology 2 (2012), pp. 2369–2392.

[27] S. Vijayavenkataraman, L. Zhang, S. Zhang, J. Y. H. Fuh, and W. F. Lu. Triply periodic minimal surfacessheet scaffolds for tissue engineering applications: An optimization approach toward biomimetic scaffold besign. ACS Appl. Bio Mater. 1 (2018), pp. 259–269.

[28] K. Wolf and P. Friedl. Extracellular matrix determinants of proteolytic and non-proteolytic cell migration. Trends in Cell Biology 21 (2011), pp. 736–744.

[29] K. Wolf and P. Friedl. Molecular mechanisms of cancer cell invasion and plasticity. British Journal of Dermatology 154 (2006), pp. 11–15.

[30] E. Yang, M. Leary, B. Lozanovski, D. Downing, M. Mazur, A. Sarker, A. Khorasani, A. Jones, T. Maconachie, S. Bateman, et al. Effect of geometry on the mechanical properties of Ti-6Al-4V Gyroid structures fabricated via SLM: A numerical study. Materials & Design 184 (2019), p. 108165.

[31] L. Yuan, S. Ding, and C. Wen. Additive manufacturing technology for porous metal implant applications and triple minimal surface structures: A review. Bioactive materials 4 (2019), pp. 56–70.

[32] M. H. Zaman, L. M. Trapani, A. L. Sieminski, D. Mackellar, H. Gong, R. D. Kamm, A. Wells, D. A. Lauffenburger, and P. Matsudaira. Migration of tumor cells in 3D matrices is governed by matrix stiffness along with cell-matrix adhesion and proteolysis. Proceedings of the National Academy of Sciences of the USA 103 (2006), pp. 10889–10894.

